# Spontaneous differentiation leads to emergence of hybrid cell states relate to poor prognosis in oral cancer

**DOI:** 10.1101/2021.08.24.457509

**Authors:** Kavya Vipparthi, Kishore Hari, Priyanka Chakraborty, Subhashis Ghosh, Ankit Kumar Patel, Arnab Ghosh, Nidhan Kumar Biswas, Rajeev Sharan, Pattatheyil Arun, Mohit Kumar Jolly, Sandeep Singh

## Abstract

**Purpose:** Cellular dynamics between phenotypically heterogeneous subpopulations of cancer cells within individual tumor is shown to be responsible for drug tolerance and overall poor prognosis; however, evidences were largely missing in oral cancer. Therefore, this study was undertaken to describe the dynamic phenotypic states among oral cancer cells, its influence on transcriptomic heterogeneity as well as its clinical significance.

**Experimental Design:** We multiplexed phenotypic markers of putative oral-stem-like cancer cells (SLCCs) and characterized diversity among CD44-positive oral cancer cell subpopulations with respect to distinct expression of CD24 and aldehyde dehydrogenase (ALDH)-activity in multiple cell lines. Population trajectories were characterized by Markov model and cell states were defined based on the population specific RNA sequencing (RNAseq). ssGSEA based gene expression signatures were explore for prognostic significance.

**Results:** Oral cancer cells followed two distinct patterns of spontaneous repopulation dynamics with stochastic inter-conversions on ‘ALDH-axis’, however a strict non-interconvertible transition on ‘CD24-axis’. Interestingly, plastic ‘ALDH-axis’ was harnessed to enrich ALDH^High^ subpopulations in response to Cisplatin treatment, to adapt a drug tolerant state. Phenotype-specific RNAseq results suggested the organization of subpopulations into hierarchical structure with possible maintenance of intermediate states of stemness within the differentiating oral cancer cells. Further, survival analysis with each subpopulation-specific gene signature strongly suggested that the cell-state dynamics may act as possible mechanism to drive ITH, resulting in poor prognosis in patient.

**Conclusions:** Our results emphasized the prognostic power of the population dynamics in oral cancer. Importantly, we have described the phenotypic-composition of heterogeneous subpopulations critical for global tumor behaviour in oral cancer; which is a prerequisite knowledge important for precision treatment, however largely lacking for most solid tumors.

**Graphical Abstract:** We have characterized diversity among CD44-positive oral cancer cells lines with respect to distinct expression of CD24 and ALDH-activity. Subpopulations showed stochastic inter-conversions on ALDH-axis but a strict non-interconvertible transition of CD24^Low^ to CD24^High^ phenotype, even in response to chemotherapy-induced stress. RNAseq study suggested the organization of subpopulations into hierarchical structure with possible maintenance of intermediate alternate states of stemness within the differentiating oral cancer cells. The described population dynamics demonstrtaed influence tumor behaviour possibly by increasing intratumoral heterogeneity in aggressive oral tumors.

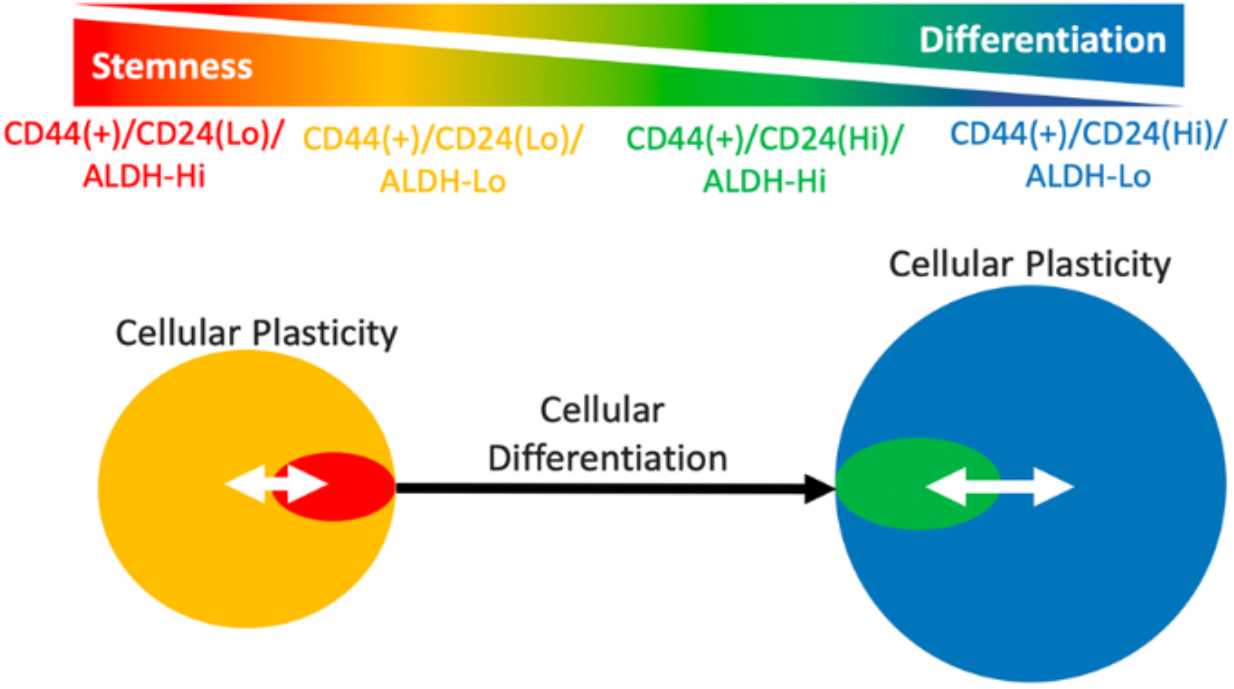

**Translational relevance:** Intratumoral heterogeneity (ITH) has been the clinically important factor, impacting aggressive cancer behaviour, drug tolerance and overall poor prognosis. Recent high-throughput studies have provided better cellular and molecular resolution of ITH; however, the prerequisite knowledge which defines the composition of subpopulations critical for global tumor behaviour is majorly lacking for most of the solid tumors. By combining phenotypic markers, we have defined four subpopulations of oral cancer cells. These subpopulations showed stochastic inter-conversions as well as a strict non-interconvertible transition among them to acheive heterogeneity. Importantly, transcriptional states of each subpopulations indicated a clinically relevant signatures for patient prognosis. Also, we observed interconversions of these subpopulations in response to Cisplatin to accumulate drug-tolerant cell state, as rapid and reversible strategy to respond to chemotherapy induced stress. Thus, the characteristics of described phenotypic subgroups may be translated to the clinic for estimating the extent of intratumoral heterogeneity in oral cancer patients.

## Introduction

Though genetic heterogeneity undeniably confers cell growth and survival advantages, it alone is not sufficient to explain several aspects of the cancer biology including functional and phenotypic heterogeneity, emergence of drug resistance and metastasis phenotypes [1]. RNA sequencing studies from multiple tumor-types showed that individual tumors with similar genetic profile may exhibit transcriptomic heterogeneity [2], that may arise due to multiple phenotypic cell states and their diverse interactions with tumor microenvironment [3–5]. Recent studies are uncovering the presence of extensive non-genetic heterogeneity at phenotypic and functional levels within genetically identical cancer sub-clones [6–8].

Ability of genetically identical cells to exist and switch to multiple phenotypic states is a long-standing notion of developmental biology [9]. However, cell state dynamics in context of generating phenotypic heterogeneity in cancer cell populations, independent of their genetic background has garnered attention, recently [4]. Studies showed that the cell state transitions can either occur spontaneously, or due to perturbations driven by external triggers [10–12]. Cells existing in each phenotypic state may exert distinct functions, such as therapy resistance, that overall benefits the tumor [13]. Commonly studied cellular processes associated with phenotypic state transitions are epithelial to mesenchymal transitions (EMT) and cancer stemness, which maybe inter-related [14]. Stem-like cancer cells (SLCCs) contribute to intratumoral heterogeneity by giving rise to different cellular phenotypes through self-renewal and differentiation [15].

Multiple studies proved the existence of SLCCs in solid tumors [16]; however, phenotypic diversity among these cells has been a major challenge in accurately defining these subpopulations in solid tumors [17, 18]. Furthering this complexity, recent evidence suggests that rather than being fixed cellular entities, stem-like characteristics can be achieved over time, even by differentiated cells [19, 20]. Hence, the unidirectional rigid hierarchy of SLCC models is widely debated [21, 22]. Thus, cancer stemness provides a conceptual framework for interrogating the aspects of complex cellular hierarchy, functional heterogeneity and plasticity in cancer.

Tumor cells with high-CD44 expression were the first identified SLCCs in Head and Neck Squamous Cell Carcinoma (HNSCC) [23]. Later, cells with combined expression for high-CD44 and aldehyde dehydrogenase (ALDH) enzyme, i.e., CD44^+Ve^/ALDH^High^ cells, were shown to enrich for HNSCC-SLCCs [24]. Simultaneously, cells with CD24^Low^/CD44^High^ cell surface marker profile were reported as representatives of SLCCs and EMT phenotypes in oral cancers [25]. Co-existence of heterogeneous oral-SLCCs with respect to EMT and their spontaneous interconversions has also been reported [26]. Despite these qualitative demonstrations, studies on population-dynamics in a quantitative and unbiased manner are largely missing in oral cancer. Questions like, at what reproducibility these subpopulations arise or relation between diverse subpopulations with respect to molecular states, has remained unanswered in oral cancer.

Here, we have attempted to investigate the cellular dynamics among oral cancer subpopulations derived by multiplexing phenotypic markers of putative oral-SLCCs, CD44, CD24 and ALDH-activity, and showed that distinct cellular phenotypes co-exist in multiple cell lines. While subpopulations with differential ALDH-activity showed stochastic inter-conversions, those with differential CD24 expression followed strict developmental hierarchy, even in response to chemotherapy-induced stress. We further identified transcriptional states of different subpopulations and showed that loss of transcriptomic signature specific to the reported putative oral-SLCCs subpopulation, suggestive of more heterogeneous tumor, was significantly associated with poorer prognosis of TCGA HNSCC patient cohorts. Overall, our work has provided evidence of cellular dynamics leading to transcriptomic heterogeneity and its influence on tumor behaviour in oral cancer patient.

## Materials and methods

### Cell culture

Oral cancer cell lines, GBC02 and GBC035 used in this study were recently established by us and maintained as described [44]. Other oral cancer cell lines SCC-029, SCC-070, SCC-084 and SCC-032 were kindly provided by Dr. Susanne M. Gollin, University of Pittusburgh, USA [45]. These cell lines were maintained in MEM (Minimum Essential Media, Cat #11095-080) with non-essential amino acids (Cat # 11140-050), L-glutamine (Cat # 25030-081) and 10% FBS.

### Dissociation of primary oral tumor cells

This study (EC/GOVT/01/12) was approved by the Institutional ethics committee of National Institute of Biomedical Genomics (NIBMG) and the institutional review board of Tata Medical Center (TMC), Kolkata, India. Primary tumor tissue samples were collected and processed as previously described by us [44, 46]. Following enzymatic digestion of tumor tissues using 1x Collagenase/Hyaluronidase and DNase-1 (Cat # 07912 & 07900; Stem cell technologies), dissociated tumor cells were collected in a different tube and filtered through 40 µ strainer. Ice-cold serum-free growth media was used to rinse the strainer’s membrane to avoid cell loss. Next, cell suspension was centrifuged at 500g for 5 minutes and supernatant was discarded. Pellet was resuspended in ACK lysis buffer (Cat # A10492-01; Gibco) for 1 minute at room temperature to lyse RBCs. It is followed by a quick centrifugation for 2 minutes, discarding of the lysis buffer and two washes with ice-cold HBSS buffer (Cat # 14175; Thermo) containing 0.5% FBS and 10mM HEPES (Cat # 15630, Thermo) (HBSS+). Cells were finally resuspended in ice-cold HBSS+ buffer and counted using trypan blue on a hemocytometer. For downstream flow cytometry analysis, dissociated tumor cells were Fc receptor blocked by incubating 1*10^6^ cells/50-100 µl of staining buffer with 2.5 µg of Human Fc Block (Cat # 564220; BD) for 10 minutes.

### Flow cytometry for combined ALDEFLUOR assay and antibody immunophenotyping

Single cell suspensions were prepared by resuspending 1*10^6^ cells /ml (for cell lines) or 1*10^6^ cells/100 µl (for dissociated primary tumor cells) of Aldefluor assay buffer. Aldefluor assay was performed as per the manufacturer’s protocol (Cat. # 01700, Stem Cell Technologies). For negative control, Diethyl aminobenzaldehyde (DEAB), an inhibitor of Aldehyde dehydrogenase enzymes, was used. Few cells (for instance; 2*10^5^ cells in 200 µl assay buffer) were separated in a tube labeled ‘with DEAB’ and 1 µl of DEAB reagent per 100 µl of sample was added and incubated for 5 minutes at 37^0^C. Next, Aldefluor reagent, BODIPY-aminoacetaldehyde (BAAA) was added in both ‘with DEAB’ & ‘without DEAB’ tubes @ 5µl/ml of assay buffer. They were incubated for 1 hour at 37^0^C, by intermittent mixing and inverting of the tubes every 15 minutes. After 1 hour, cells were washed with ice-cold Aldefluor assay buffer for 5 minutes at 400 rcf and supernatant is discarded. Further, ‘with DEAB’ control tube is sub-divided into two tubes labeled ‘with DEAB only’ and ‘with DEAB+Antibody’. CD24-Alexa647 (Cat # 561644; Clone: ML5; BD) and CD44-BV786 (Cat # 564942; Clone: G44-26; BD) antibodies were added in ‘without DEAB’ and ‘with DEAB+Antibody’ tubes and incubated for 30 mins at 4^0^C. In case of primary tumor cells, hematopoietic lineage cocktail antibodies (Cat # 22-7776-72; eBioscience) were added along with CD24-BV421 (Cat #562789; BD) and CD44-BV786 (Cat #564942; BD) antibodies. After incubation, cells are washed twice with ice-cold Aldefluor buffer, filtered and resuspended @ 3*10^6^ cells/ml. Propidium iodide (Cat # P4170; Sigma) at final concentration of 2µg/mL was added to exclude dead cells from the analysis. Flow cytometry data was acquired using BD FACS Aria Fusion cytometer and analysis was performed using FCS Express 5 (DeNovo Software).

### Cell Sorting

Sorting buffer was prepared by adding 5% FBS and 1X antibiotic-antimycotic mix (Cat # 15240-062, 50-0640; Thermo) to HBSS+ buffer. FACS tubes with sorting buffer were kept at 4^0^C, 2 hours prior to cell sorting. Gating was set as per the experimental choice and sort layouts were prepared. For isolating the four subpopulations, cells were sorted four ways into the collection tubes containing sorting buffer, using BD FACS Aria Fusion flow cytometer.

### Analysis of frequencies of four subpopulations

Percentages of four phenotypic subpopulations were obtained from the gating statistics of flow cytometry dot plots, analyzed by FCS Express 5 Software. Percentages were summed up to 100% and plotted as stacked bar graphs, using graphpad prism software.

### Cell Trace (CT)-Violet dye dilution assay combined with marker phenotyping

Assay was performed as per the protocol described previously [44], with minor modifications. Equal numbers of parental cells were seeded in triplicates (1X10^5^ cells per 60mm dish). 48 hours later CT-Violet dye staining (Cat # C34557, Invitrogen) was performed, followed by Aldefluor and CD24 immuno-staining on one of the stained plates, for recording baseline fluorescence signal at Day 0. Other plates were allowed to proliferate for 4 days and subsequently subjected to Aldefluor and CD24 immuno-staining. Proliferation index was calculated using FCS express software based on fluorescence intensities at Day-0 and Day-4. Readings from three independent plates were taken to determine average proliferation index and SD.

### Repopulation Assay

Sorted cells were centrifuged at 400 rcf for 5 minutes. Supernatant is discarded and cells are resuspended in their respective growth media and plated in wells of a 24-well culture plate @ 5000 cells/well in triplicates. Cells are allowed to grow un-interrupted for 10 days with media change once in every three days. On 10^th^ day post-sort, cells were harvested using Accutase dissociation reagent and re-analyzed for Aldefluor assay, CD24 and CD44 immuno-phenotyping as described above.

### Cisplatin treatment and Repopulation assay

After plating cells for repopulation, on 6^th^ day post-sort, growth media containing 2µM Cisplatin was added to cells. Following 48 hours of treatment (i.e, 8^th^ day post-sort), Cisplatin was removed and cells were recovered by adding fresh growth media devoid of Cisplatin, for next 48 hours. On 10^th^ day post-sort, cells were harvested and re-analyzed for Aldefluor assay, CD24 and CD44 immuno-phenotyping as described above.

### Spheroid Formation Assay

As previously described [46], 5-10 cells/ul were resuspended in growth media without serum and with 20ng/ml EGF (Thermo), bFGF (Thermo), 1x B-27 (Thermo) and 0.4ug/ml hydrocortisone (Merck, Sigma) along with 1.25% geltrex (Thermo) and plated in multiple wells of ultra-low attachment plate (Corning). Growth factors were supplemented every alternate day. Spheroids formation was continued for 7-10 days. Spheroids with size ≥60 µ were counted for calculating efficiency. Size of spheroids was counted using Image J software. Limiting dilution assays were conducted in same manner with varying cell numbers ranging from 5-500/well, as depicted in results section. For estimation of SLCCs frequencies in each subpopulation, ELDA (Extreme Limiting Dilution Analysis) was performed by entering the limiting dilution spheroid data in the website http://bioinf.wehi.edu.au/software/elda/.

Primary spheroids generated were collected and centrifuged down at 800 rcf for 5 minutes. Supernatant was discarded and to the pellet, Dispase (Cat # 07923; Stem cell technologies), Collagenase (Cat # & 07909; Stem cell technologies) mix in 1:1 ratio was added to digest geltrex by incubating for 10-15 minutes at 37^0^C. TNS was added to stop over-digestion and centrifuged for 5 minutes at 800 rcf. Spheroids were further suspended in Trypsin-EDTA for 2-5 minutes to obtain single cells. Reaction was stopped by another round of TNS addition and removal of supernatant. Finally, single cells were resuspended in fresh, serum free, spheroid formation media described above and gently pipetted for homogeneity. They were plated for secondary spheroid formation at the concentrations of 5-500 cells/well.

### 3D Repopulation Assay

Sorted cells were plated for spheroid formation (5 cells/µl) as described above and their growth was followed for 7-10 days. Single spheroids >60 µ size from each subpopulation were carefully picked by observing under the microscope and plated in wells of a 96 well plate with one spheroid per well. They were cross checked microscopically and wells with only single spheroids were followed up. Once colonies grew, they were transferred onto dishes with a bigger surface and the repopulation analysis was performed from 4 independent single spheroid colonies from each subpopulation.

### RNA Isolation, qPCR

RNA isolation and qPCR were performed as described previously [44], using RNeasy mini/micro kit and SsoEva Green SYBR mix on CFX96 Real-Time PCR system (BioRad) respectively. Relative gene expression fold changes were calculated by 2^−ΔΔCT^ method. Sequences of the primers used are provided as table below.

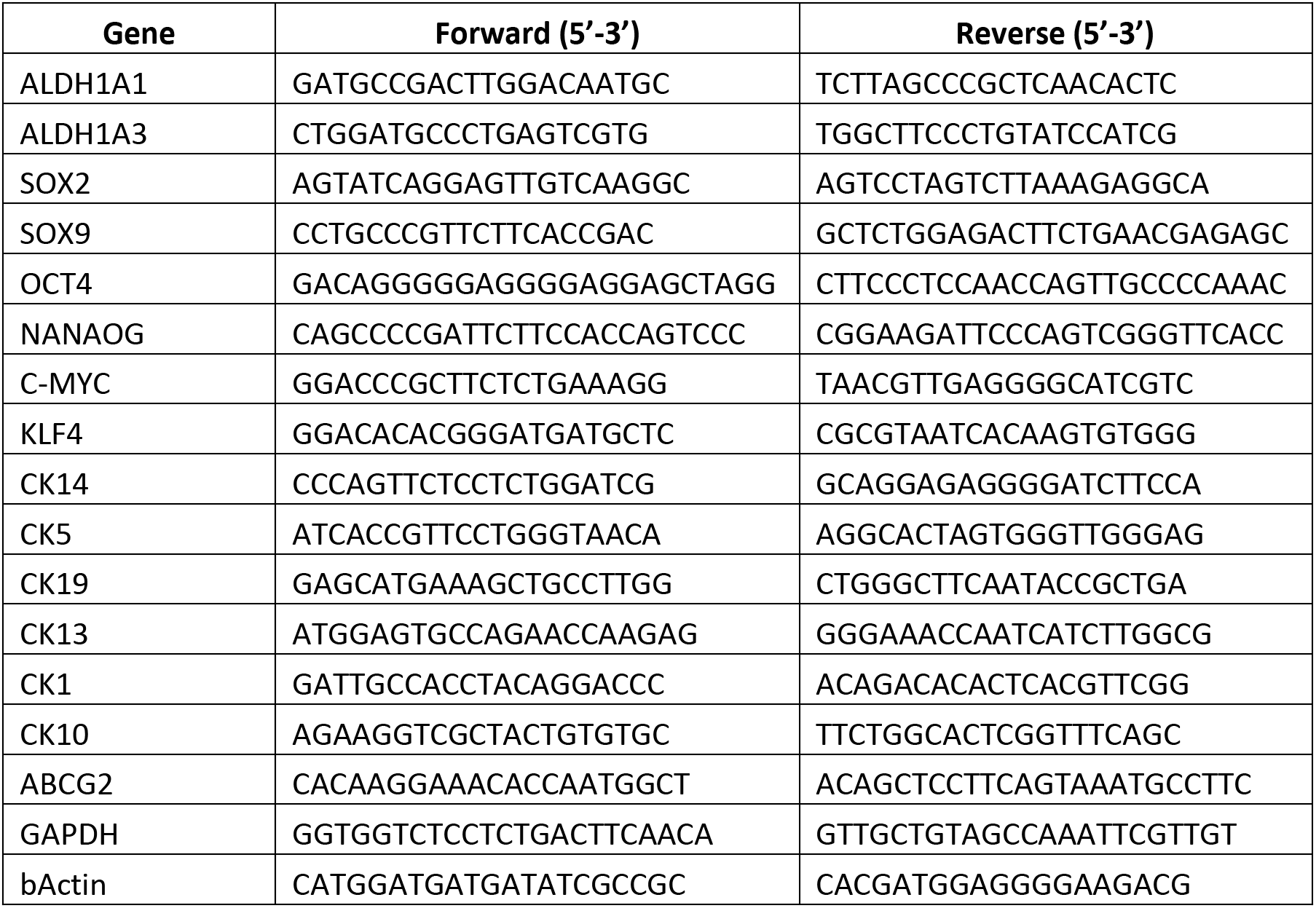

### Discrete time markov chain model construction

A discrete time markov chain model was constructed to explain the repopulation patterns of different phenotypes. Assumptions of the model are as follows:

1. Repopulation is due to transition of different phenotypes between each other. Other processes like cell division do not affect the phenotypic heterogeneity.
2. Transition rates among the phenotypes are independent of time and of each other.
3. infer the transition rates, the package *CellTrans* [27](R 3.6.4) was applied on repopulation data. Briefly, the package uses repopulation data at time *t_i_* obtained starting from a sorted population to construct a phenotypic proportion matrix (*PPM_ti_*). Using the theory of Markov chains, it calculates the transition matrix (*TM*) corresponding to the unit time by left-multiplying the proportion matrix with the identity matrix and taking a *t_i_^th^*root of the resultant matrix.

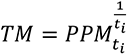

The repopulation trajectories in Fig 4 were constructed using the transition matrix using the following formula:

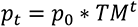

Where *p*_0_ is the population composition (a row vector) at time *t* = 0 and *p_t_* is the population composition at time *t* = *t*.

For sensitivity analysis of the estimated transition matrices, we generated random transition matrices by randomly shuffling the rows of the estimated TM and then randomly shuffling the columns.

### Transition matrix randomization

From the markovian model, we were able to obtain transition matrices and predicted possible trajectories leading to steady states. To understand our predictions further and to verify whether the predicted trajectories are unique to the transition matrix and hence to the experimental data, we randomized transition matrices and compared the matrices and the steady states with those of “WT” matrix. To maintain transition matrix property, i.e., all row-sums must be equal to one, the randomization was performed by shuffling rates in a single row and shuffling entire rows (Supplementary Fig 7E).

We measured the distance between steady states and the element-wise distance between the transition matrices using a standard distance metric: Euclidean distance. Based on these distances, we clustered the transition matrix-steady state pairs using kmeans function of R, as demonstrated for the case of GBC02 in Supplementary Fig 7E. Further analysis of these clusters in terms of mean and standard deviation values of steady states (Supplementary Fig 7G-H) and transition rates (Supplementary Fig 7 I-J) were performed.

### RNA sequence data generation, analysis, and identification of differentially expressed genes

For RNA sequencing studies, RNA is extracted from FACS isolated four subpopulations of the GBC02 cell line grown in 2D monolayer cultures, in triplicates, and also from the ‘Red’ and ‘Green’ spheroids generated from sorted cells of GBC02 cell line, in duplicates.

Whole RNA was extracted from three replicates for each of the cell-lines followed by a quality assessment using Agilent Bioanalyser-2100 with Agilent nano Kit (Agilent Technologies). For each sample, library was prepared by taking 400ng of RNA and sequencing was performed using Illumina Hiseq-2500 to obtain 50 million paired end reads. Raw data QC was performed through FastQC, and the RNA sequence data was aligned to hg19 primary assembly using STAR aligner (version 2.6.0) with GRCh37.87 (Ensembl) transcript model. An average of 87% uniquely mapped reads was obtained from each sample. HTSeq (version 0.11.0) was used to generate count data. Protein coding genes that expressed in at least two of three biological replicates in case or control group were taken for differential gene expression (DGE) analysis. DGE analysis was performed through DESeq2 method with the raw count data. A gene was considered to be differentially expressed if |log2 fold change| > 1 with FDR corrected p-value < 0.1 is obtained. All downstream analysis was performed in R-3.5.1 and ggplot2 package was used for generation of plots.

### Extraction of cell population-specific unique DEGs

Unique DEGs were obtained for each of the cell subpopulations based on all the pairwise comparisons. Genes were selected as unique DEGs for a cell population *i*as following - (1) by taking common set of n different set of DEGs (all pairwise comparison of cell population *i* with other cell population). (2) by excluding genes from common sets that are differentially expressed for any other pairwise comparison.

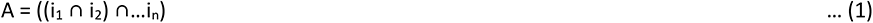

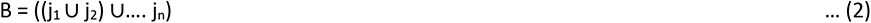

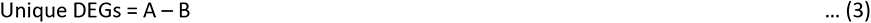

Here, i_1_ to i_n_ are set of DEGs of cell population i as compared to other n numbers of different cell populations and j_1_ to j_n_ are set of DEGs for all other cell population pairwise comparisons.

### Calculation of population-specific fold change

Cell population-specific fold change for each unique DEG for a given cell population was calculated as follows:

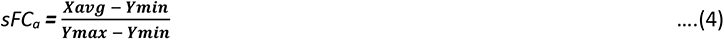

sFCa is specific fold change of gene “a” in X cell population .Where,

X_avg_ – Mean expression of gene “a” in the X cell population

Y_min_– Minimum of average expression of gene “a” in other cell populations

Y_max_– Maximum of average expression of gene “a” in other cell populations

### ssGSEA scores and survival analysis utilizing population-specific signature

The scores were calculated using GSVA R Bioconductor package and “ssGSEA” method was employed in the estimation of gene-set enrichment scores per sample.

To identify the role of different cell population gene signatures on patient survival, Head and Neck Cancer patients were classified into high and low enrichment groups, using the mean enrichment value as cut-off of single sample gene set enrichment analysis (ssGSEA) scores of a gene signature. To perform survival analysis R tool, *survival* was used.

This analysis was performed as follows:

i. Select top n number of up and down regulated genes from unique DEG list sorted based on *specific Fold Change (sFC)* values.
ii. Calculate *ssGSEA* scores using the selected genes in the previous step.
iii. Segregate patients into “high” and “low” group. The HNSCC patients that showed *ssGSEA* scores above the mean *ssGSEA* value, were classed as the high expression group, and those below the mean value were classed as the low expression group.
iv. Perform survival analysis between the two patient groups.

## Results

### ALDH^High^ cells exhibited both CD24^Low^ and CD24^High^ phenotypes

We first examined the co-expression of markers related to putative oral-SLCCs; CD44, CD24 and ALDH in five different oral cancer cell lines. All tested cell lines were positive for CD44 marker expression. Three of five cell lines (GBC02, SCC-070 and SCC-029) showed co-existence of both CD24^High^ and CD24^Low^ sub-populations (Fig 1A, 1B and Supplementary Fig 1A, B), whereas, GBC035 and SCC-032 were of mainly CD24^High^ and CD24^Low^phenotypes, respectively (Fig 1B and Supplementary Fig 1C,D). Analysis for ALDH-activity by ALDEFLUOR-assay showed that ALDH^High^ cells were not only present in previously reported CD24^Low^ subpopulation of CD44-positive cells, but also in relatively unexplored subpopulation with CD24^High^ phenotype (Fig 1A, B and Supplementary Fig 1). Hence, based on the differential status of these two markers, we identified four sub-populations of CD44-positive cells with marker profiles, CD24^Low^/ALDH^High^, CD24^Low^/ALDH^Low^, CD24^High^/ALDH^High^, CD24^High^/ALDH^Low^ which were termed as ‘Red’, ‘Orange’, ‘Green’ and ‘Blue’ subpopulations, respectively. Frequencies of each of these subpopulations were analyzed in all tested cell lines. Importantly, ALDH^High^ phenotypes (‘Red’ and ‘Green’) were maintained at lower frequencies, while ALDH^Low^ subpopulations (‘Orange’ and ‘Blue’) were the dominant ones across different cell lines (Fig 1B). Overall, these results emphasized the co-existence of ALDH^High^ cells overlapping with both CD24^High^ and CD24^Low^ phenotypes in CD44-positive oral cancer cells. Further, phenotypes observed in cell lines were also present in primary oral tumors (gating strategy, Supplementary Fig 2). We found ALDH^High^ cells to be primarily enriched in CD24^Low^ phenotype (Fig 1C, D, pie chart, Supplementary Fig 3). However, ALDH^High^ cells were also enriched in CD24^High^ subpopulations in primary oral tumors at frequencies ranging from 0.3-11% (Fig 1C, D, Supplementary Fig 3). Collectively, results provided evidence for existence of heterogeneous ALDH^High^ cells with diverse CD24 related phenotypes in primary oral tumors similar to *in-vitro* cell lines.

**Figure 1:**
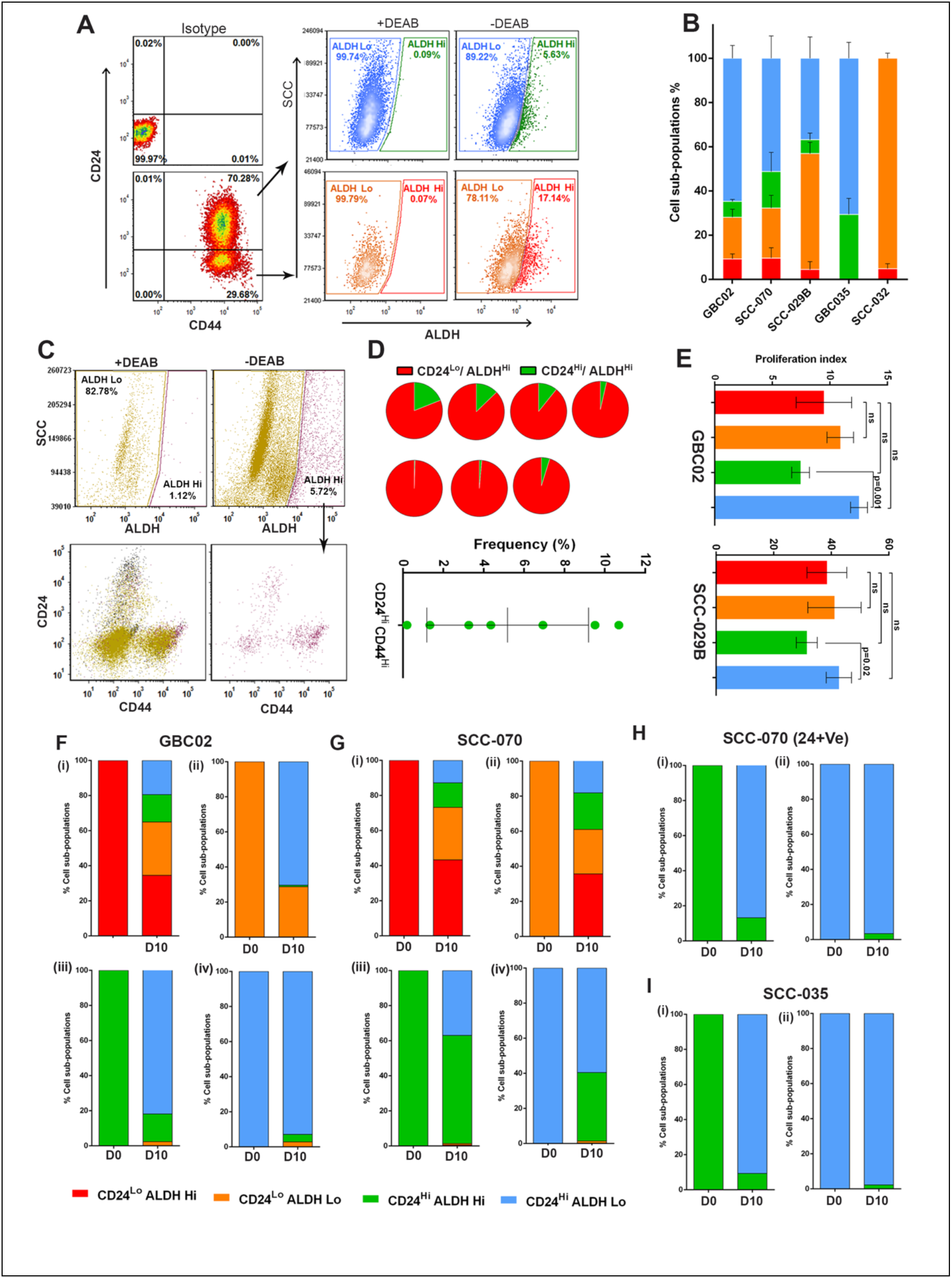
Subpopulations exhibited distinct repopulation dynamics: **(A)** Representative FACS dot plots of GBC02 cell line’s CD24/CD44 staining (Left bottom) with respect to Isotype control (Left top). ALDEFLUOR (ALDH) phenotype of CD24^High^/CD44^+Ve^ sub-population (Right top) and CD24^Low^/CD44^+Ve^ sub-population (Right bottom) in presence or absence of DEAB. **(B)** Frequency distribution of the four sub-populations CD24^Low^/ALDH^High^ (Red), CD24^Low^/ALDH^Low^ (Orange), CD24^High^/ALDH^High^ (Green) and CD24^High^/ALDH^Low^ (Blue) in five genetically distinct oral cancer cell lines. Error bars represent mean ± SEM from three biological repeats. **(C)** Representative FACS dot plots of a freshly resected and digested patient tumor.Hematopoietic lineage negative cells segregated into ALDH^High^ and ALDH^Low^ cells in presence or absence of DEAB (top). CD24/CD44 phenotype of all lineage negative cells (bottom left). CD24/CD44 phenotype of ALDH^High^ cells only (bottom right). **(D)** Pie charts showing frequencies of CD24^High^/ALDH^Hi^(Green) and CD24^Low^/ALDH^High^(Red) subpopulations from 7 freshly resectedhuman oral tumor samples and frequencies of CD24^High^/ALDH^High^cells from all 7 oral tumor patient samples. **(E)** Graphs showing proliferation index of GBC02 and SCC029 cell lines based on CellTrace-Violet dye dilution assay. **(F, G, H, I)** Repopulation frequencies of each subpopulation in (i) Red (ii) Orange (iii) Green and (iv) Blue sorted cells on Day-0 and Day-10 of sorting in GBC02 (F), SCC-070 (G), GBC035 (H) and SCC-070 (CD24+) sub-line (I).

### Subpopulations exhibited distinct repopulation dynamics

Maintenance of subpopulations at different frequencies in a mixed population could be due to differences in their proliferative potential. However, in ‘Cell Trace-Violet’ dye dilution assay, we did not observe significant differences in proliferation indices among the four subpopulations in both tested cell-lines, GBC02 and SCC-029B (Fig 1E). This observation prompted us to probe the repopulation dynamics of these four subpopulations. Repopulation results from GBC02, SCC-029B and SCC-070 cell lines showed that the ‘Red’ subpopulation was most ‘potent’ which efficiently reproduced all four subpopulations. Interestingly, ‘Green’ subpopulation could only give rise to itself and its ALDH^Low^ counterpart, showing ‘commitment’ to generate only CD24^High^ phenotype in all the tested cell lines (Fig 1F,G and Supplementary Fig 4A,B,C). While the ALDH^Low^-‘Orange’ subpopulations efficiently repopulated the ALDH^Low^ subpopulations of ‘Orange’ and ‘Blue’ phenotypes; the ‘Blue’ subpopulation showed the phenomena of ‘CD24^High^ specification’ by repopulating itself with higher efficiency (Fig 1F,G and Supplementary Figure 4A,B,C). Of note, the ALDH^Low^ (‘Orange’ or ‘Blue’) subpopulations showed higher plasticity and generated ALDH^High^ phenotypes at higher frequency in SCCC-070 cell line as compared to other cell lines (Fig 1G, Supplementary Fig 4B). Distinct from other cell lines, SCC-032 could not generate the CD24^High^ (‘Green’ or ‘Blue’) from CD24^Low^ (‘Red’ or ‘Orange’) subpopulations. Also, similar to the ‘Orange’ subpopulation of SCC-070, the ‘Orange’ subpopulations of SCC-032 demonstrated the capacity to recapitulate both ‘Red’ and ‘Orange’ subpopulations (Supplementary Fig 4D).

Further, to confirm the property of CD24^High^ cells to be a committed subpopulation, we developed a separate sub-line termed as ‘SCC-070 (CD24+)’, by sorting and propagating CD24^High^ cells repopulated from ‘Red’ subpopulation of SCC-070 cells. Interestingly, though CD24^High^ cells were originated from CD24^Low^ subpopulation, these cells did not regenerate CD24^Low^ cells or parent phenotype (Fig 1H, Supplementary Fig 5A(i)), confirming the committed property of CD24^High^ subpopulation. Further, the specification of ‘Blue’ subpopulation could also be confirmed (Fig 1H, Supplementary Fig 5A(ii)). These results were further supported by an independent patient derived cell line, GBC035, which predominantly has phenotypically homogenous CD24^High^ phenotype. Similar to SCC-070 (CD24+) subline, sorted ‘Green’ and ‘Blue’ subpopulations of GBC035 also showed ‘Green’ subpopulation to be committed to generate only CD24^High^ subpopulations; whereas ‘Blue’ subpopulation to have specification for generating itself (Fig 1, Supplementary Fig 5B).

Although with certain cell-line specific trends; overall, we uncovered two distinct patterns of repopulation dynamics; 1) ‘Fixed’ CD24-Axis, where transitions from CD24^Low^ to CD24^High^ were largely unidirectional and non-interconvertible, and 2) ‘Plastic’ ALDH-axis, where transitions from ALDH^High^ to ALDH^Low^ and vice-versa were frequent.

### All subpopulations exhibited enrichment of spheroid forming cells and maintained repopulation dynamics

Higher potency of ‘Red’ and ‘Orange’ (CD24^Low^) as compared to ‘Green’ and ‘Blue’ (CD24^High^) subpopulations indicated a possible hierarchical cellular organization, with putative oral-SLCCs to be CD24^Low^, as has been earlier reported [25]. Therefore, we next tested all four subpopulations for *in vitro* spheroid formation assays in three cell lines. However, surprisingly all four subpopulations from GBC02 cell lines demonstrated spheroid-forming potential in tested two generations of spheroid propagation (Fig 2A-i). Next, more critical evaluation of their spheroid-forming efficiencies were performed by limiting dilution spheroid formation assays. This revealed the highest enrichment of spheroid-forming cells (1 in 3 cells) in the ‘Red’ subpopulation. However; interestingly, the ‘Orange’ (1 in 6 cells), ‘Green’ (1 in 11 cells) and ‘Blue’ (1 in 24 cells) subpopulations also demonstrated the enrichment of spheroid-forming cells (Fig 2B-i; 2C-i). These differences in efficiency were significant as tested by pair-wise comparisons (Fig 2C-i, last column). Similar to these observations; SCC-070 (CD24+Ve) sub-line also demonstrated spheroid-forming efficiency of ‘Green’ (1 in 6 cells) to be higher than ‘Blue’ subpopulation (1 in 16 cells) (Fig 2A-ii 2B-ii, 2C-ii). Interconverting ‘Red’ and ‘Orange’ subpopulations of SCC032 showed similar (1 in 159 cells) spheroid-forming efficiency (Fig 2A-iii and 2B-iii, 2C-iii).

**Figure 2:**
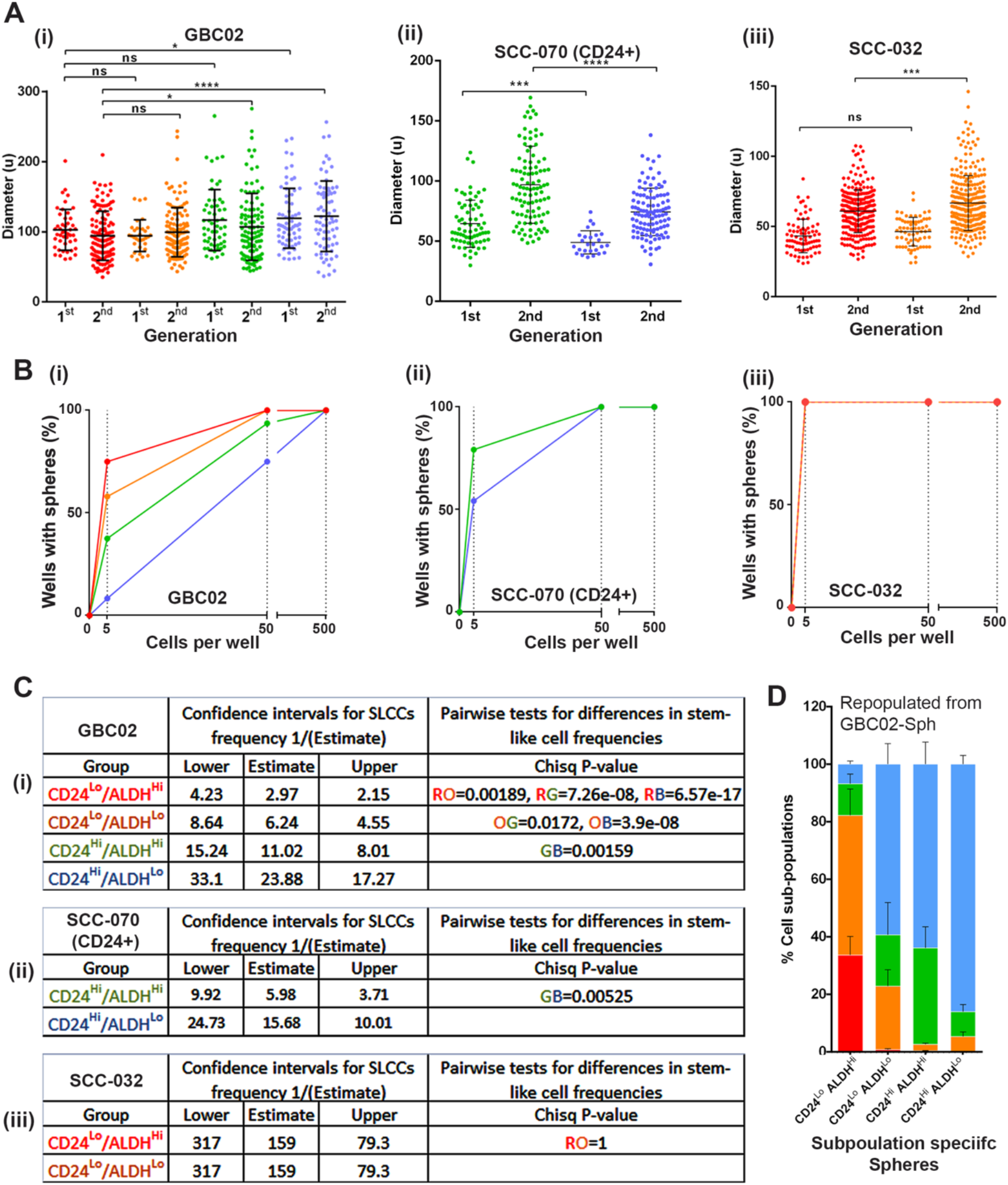
Enrichment of cells with spheroid forming ability by all four subpopulations: **(A)** Size of the four subpopulations spheroids from GBC02, SCC-070 CD24+ and SCC-032 cell lines. **(B)** Second generation spheroid formation frequencies in limiting dilution assay from GBC02, SCC-070 CD24+ and SCC-032 cell lines. **(C)** Estimation of SLCC frequencies in each subpopulation of (i) GBC02, (ii) SCC-070 CD24+ and (iii) SCC-032 cell lines using ELDA software. **(D)** Frequencies of each subpopulation in colonies generated from Red, Orange, Green and Blue subpopulation specific single spheroids after 30 days of sorting. Error bars represent mean ± SEM from four biological repeats.

We next tested if clonal spheroid-cultures of these four subpopulations exhibit similar patterns of repopulation as previously observed for 2D cultures (Figure 1F-I). Intriguingly, with close resemblance to the repopulation dynamics in 2D colonies, each clonal spheroid-cultures of these subpopulations exhibited similar pattern of repopulation (Fig 2D, Supplementary Fig 6A, B). Spheroids generated from ‘Red’ subpopulation remained the most potent to generate all four subpopulations, whereas the ‘Green’ subpopulation spheroids remained committed to generate CD24^High^ subpopulations and ‘Blue’ demonstrated its specificity to mainly maintain itself, efficiently (Fig 2D, Supplementary Fig 6B). These results suggested that, Red-subpopulation exerts its supremacy with highest spheroid forming efficiency and repopulation potency, compared to other subpopulations. However, importantly other subpopulations too showed spheroid formation potential, albeit at a lower frequency than the ‘Red’. Overall, data from all different cell lines emphasized that CD24^High^ cells possessed the ability to initiate spheroid formation despite being committed to repopulate only CD24^High^ cells.

### Quantitative model of phenotype-transition dynamics

Next, to demonstrate all possible transition paths which these subpopulations may take in terms of self-maintenance or conversions resulting in the observed repopulation dynamics, we applied a Discrete Time Markov Chain (DTMC) model for a simple linear description of the evolution of phenotypic heterogeneity based on their mutual transitions. Assuming that the transition rates are constant over time, we used *CellTrans* [27] package to infer daily transition rates of all four subpopulation among each other in GBC02, SCC-029B, SCC-070, GBC035 and SCC032 cell lines. Results were represented as transition graphs (Fig. 3A-i, ii, iii, iv). All subpopulations showed high sustenance rates (rate with which the cells maintain their identity), with ‘Blue’ subpopulations having the maximum. Also, majority of transition paths led to the ‘Blue’ subpopulation with very low transition rate from ‘Blue’ subpopulation to other subpopulations (efflux rate). Thus, quantitative modelling explained the experimental observation of low efficiency of transition to other subpopulations from sorted ‘Blue’ subpopulation.

**Figure 3:**
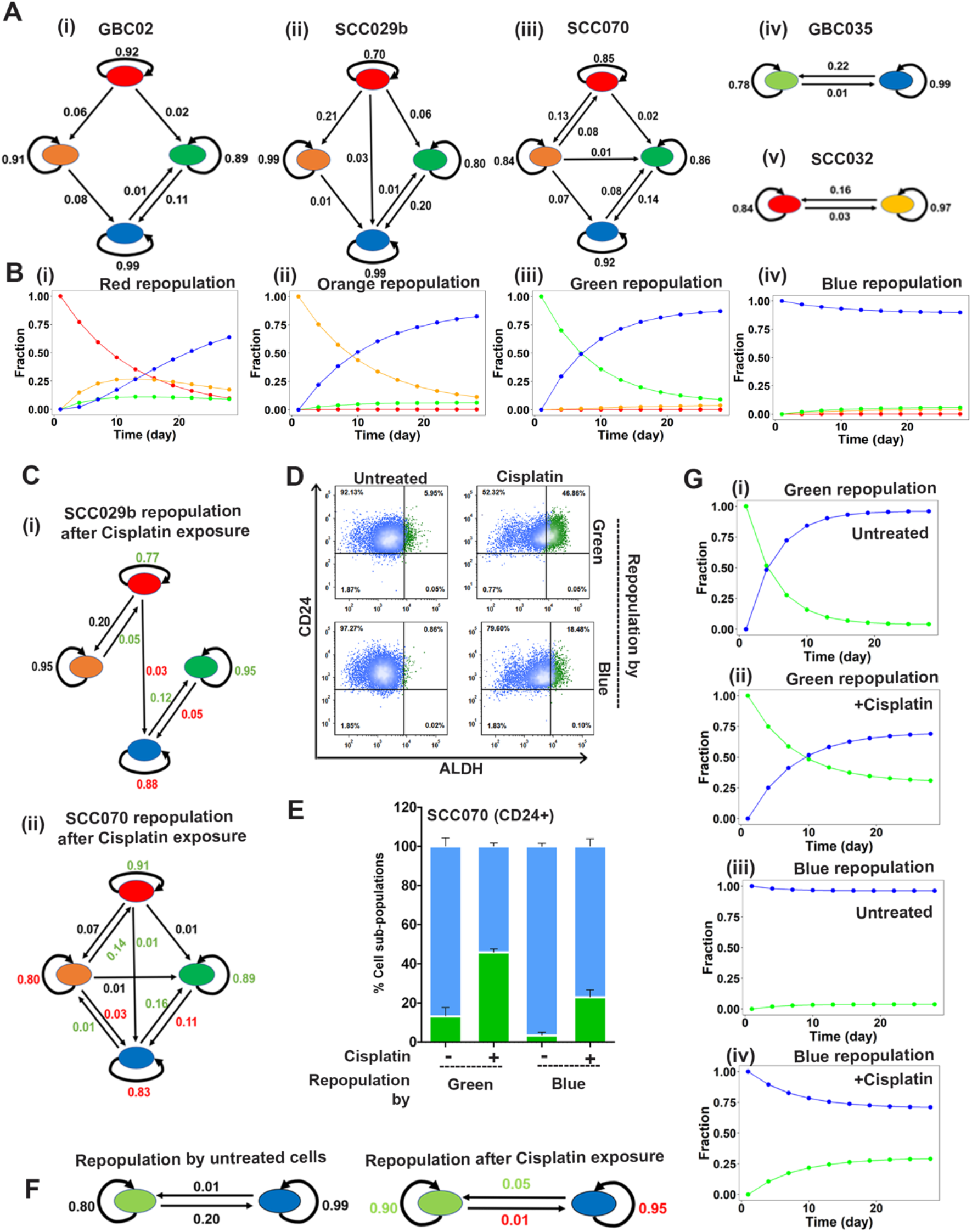
Spontaneous and Cisplatin induced population dynamics: **(A)** The phenotypic transition graphs of (i) GBC02, (ii) SCC029, (iii) SCC070, (iv) GBC035 and (v) SCC032. Arrows represent the direction of transition and the corresponding labels show the calculated transition rates. Lack of arrows between 2 subpopulations indicates that the transition rate in that direction is less than 0.01. **(B)** The phenotypic transition trajectories predicted based on the transition graph for GBC02 cell line’s (i) Red (ii) Orange (iii) Green (iv) Blue subpopulations repopulations.**(C)** The phenotypic transition graphs of SCC029 and SCC070 subpopulations treated with 2µM Cisplatin showing increased influx and sustenance rate of the green state and its reduced outflux, overall increasing the fraction of green subpopulation under Cisplatin treatment. **(D)**Representative dot plots of SCC070 CD24+ sub-line’s repopulation showing increase in Green cells in sorted Green (top) and Blue (bottom) subpopulations in untreated and 2µm Cisplatin treated conditions. **(E)** Repopulation frequencies of SCC070 CD24+ sub-line’s Green and Blue subpopulations in untreated vs. Cisplatin (2µm) treated conditions. Error bars represent mean ± SEM from two biological repeats. **(F)** The phenotypic transition graph of Green and Blue subpopulations of SCC070 (CD24+) sub-line in untreated and Cisplatin treated conditions. **(G)** The phenotypic transition trajectories of (i, iii) Untreated Green and Blue cells and (ii, iv) Cisplatin treated Green and Blue cells repopulations of SCC070 CD24+ sub-line showing increased fraction of Green cells in the population upon Cisplatin treatment.

Using these calculated transition rates, we next generated trajectories of evolution of population heterogeneity starting from sorted homogeneous subpopulations using the calculated transition rates. As shown in Figure 3B for GBC02 cell line, when starting from a ‘Red’ sorted subpopulation, reduction in the fraction of ‘Red’ subpopulation was associated with an initial rise and fall in the ‘Orange’ subpopulation and rise in ‘Blue’ subpopulation, indicating a transition from CD24^Low^ to CD24^High^ phenotype (Fig 3B-i). Furthermore, initial population of homogeneous CD24^High^ phenotypes (‘Blue’ or ‘Green’ subpopulations) failed to give rise to CD24^Low^ phenotypes, while the homogeneous ‘Green’ subpopulations CD44^+^/CD24^High^/ALDH^High^ could give rise to CD44^+^/CD24^High^/ALDH^Low^ ‘Blue’ subpopulation and vice-versa (Fig 3B-iii, iv). These trajectories revealed the direction(s) of population dynamics along the ‘CD24 and ALDH-axes’. Together, these results, obtained across different cell lines (Supplementary Fig 7A-D) reaffirm our experimental observations of uni-directional transition in ‘CD24-axis’ and bi-directional plasticity in ‘ALDH-axis’.

Next, we performed sensitivity analysis to characterize the effects of cell-to-cell variability in transition rates on emergence of observed phenotypic compositions. To do so, we shuffled the values in the transition matrices (that contain calculated transition rates), i.e. these *in silico* experiments created ‘hypothetical’ scenarios where, for instance, the influx and efflux rates were swapped and the effect on phenotypic distributions was calculated (see methods section for more details) (Supplementary Fig 7E-N). We found that as long as high influx and low efflux of the ‘Blue’ subpopulation rates were maintained, blue population retained its dominance. These observations identified the necessary and sufficient conditions enabling the ‘Blue’ subpopulation to be more dominant than the red, orange and green.

### Cisplatin alters phenotype-transition dynamics due to plasticity in ALDH-Axis but not CD24-axis

Previous studies showed that external cues such as chemotherapy-induced stress propels cells to transit into a more drug tolerant state [13, 28] often associated with stem-like phenotypes. Therefore, we explored the influence of chemotherapy on transition dynamics of the subpopulations for SCC-029B and SCC-070 cell lines. Exposure to Cisplatin during repopulation, resulted in ‘Red’ subpopulation retaining higher frequency of itself and generating significantly higher frequency of ‘Green’ subpopulations as compared to the ‘Red’ subpopulation unexposed to Cisplatin, for both cell lines (Supplementary Fig 8, 9). Similarly, isolated ‘Green’ subpopulation exposed to Cisplatin during repopulation retained itself at a significantly higher frequency as compared to untreated in both cell lines (Supplementary Fig 8, 9). Moreover, the ALDH^Low^ (‘Orange’ and ‘Blue’) subpopulations showed significant increase in ALDH^High^ subpopulations (‘Red’ and ‘Green’ respectively) when exposed to Cisplatin compared to untreated cells (Supplementary Fig 8, 9). Therefore, our data suggested that the Cisplatin treatment enriches ALDH^High^ subpopulations which maybe a result of conversion from ALDH^Low^ subpopulations, possibly due to their intrinsic plasticity. Furthermore, it is important to note that even under drug induced stress, CD24^High^ subpopulation retained its commitment and did not generate CD24^Low^ subpopulations.

Using the quantitative approach described earlier, we compared the transition rates in response to Cisplatin treatment with those in repopulation of treatment *naive* subpopulations. Cisplatin treatment increased the sustenance rates for both ALDH^High^ subpopulations (‘Red’ and ‘Green’) and reduced the efflux rate from ‘Red’ or ‘Green’ subpopulation to any other subpopulations in both SCC-029B and SCC-070 cell lines (Fig 3C-i, ii). Further supporting the transition of ALDH^Low^ subpopulations to ALDH^High^ subpopulations upon Cisplatin treatment, repopulation from the ‘Orange’ and ‘Blue’ subpopulations in both SCC-029B and SCC-070 cell lines were in stark contrast to observations in untreated conditions. The quantitative model demonstrated a higher rate of transition from ‘Orange’ to ‘Red’ and ‘Blue’ to ‘Green’ subpopulations which was either absent or at a lower rate in untreated cells (Supplementary Fig 10).

To confirm that the bidirectional plasticity of ALDH-Axis is harnessed by ALDH^Low^ cells to accumulate ALDH^High^ subpopulation but the CD24-axis remains unidirectional even in response to Cisplatin, we next treated Green and Blue subpopulations of SCC-070 (CD24+Ve) sub-line. As anticipated, the ‘Green’ subpopulation maintained itself at higher frequency and ‘Blue’ subpopulation repopulated higher frequency of ‘Green’ cells with Cisplatin. Importantly, both ‘Blue’ or ‘Green’ subpopulations failed to give rise to CD24^Low^ phenotypes even in response to Cisplatin treatment (Fig 3D, E). Assessment of this population dynamics revealed reduced sustenance and increased efflux rates of ‘Blue’ subpopulation, but increased sustenance rate of the ‘Green’ subpopulations in Cisplatin treated conditions (Fig 3F). Dynamic trajectories plotted to predict the evolution of subpopulations under untreated and Cisplatin treated conditions predicted that, Cisplatin treated ‘Green’ subpopulation maintained itself at higher fraction, while ‘Blue’ subpopulation generated ‘Green’ subpopulation at higher rates (Fig 3G-ii, iv) as compared to their untreated counterparts (Fig 3G-i, iii). These results strongly demonstrated the plastic nature of ALDH-Axis to be responsible for accumulation of ALDH^High^ subpopulations under Cisplatin treatment condition; however unidirectional transitions along the CD24-Axis (CD24^Low^ to CD24^High^) were strictly maintained.

### Transcriptome profiling revealed different levels of cell maturation among subpopulations

The observed dynamic relationship among subpopulations prompted us to determine the molecular association between them. Towards this, we performed RNA sequencing of these four subpopulations isolated from 2D cultures in three biological repeats and 3D-spheriods generated from ‘Red’ and ‘green’ subpopulations of GBC02. Based on the overlap between sets of differentially expressed genes (DEGs) obtained from pair-wise comparisons we found varying degrees of overlap in DEGs between different comparisons (Fig 4A). For instance, 79% of overlap was seen for DEGs between ‘Orange’ vs. ‘Blue’ subpopulations and ‘Orange’ vs. ‘Green’ subpopulations. Similarly, 76% of DEGs for ‘Red’ vs. ‘Green’ subpopulations were found in ‘Red’ vs. ‘Blue’ DEG analysis. Conversely, 71% of DEGs noted in ‘Red’ and ‘Blue’ were witnessed in ‘Red’ vs. ‘Green’ DEGs-list. (Fig 4A). This pattern of overlap pointed out that ‘Red’ subpopulation may be approximately equidistant from ‘Blue’ and ‘Green’ subpopulations. Similarly, ‘Orange’ may also be approximately equidistant from ‘Green’ and ‘Blue’ subpopulations. Further, the DEGs between ‘Blue’ and ‘Green’ subpopulations had very few genes in common when overlapped with any of the five other DEG lists (see column 1 in Fig 4A), indicating that these subpopulations may be transcriptionally most similar to one another. These overlaps in DEGs, coupled with quantifying distances among these comparisons enabled us in arranging these subpopulations in a hierarchical structure with ‘Red’ subpopulations being the most upstream, followed by ‘Orange’, ‘Green’ and ‘Blue’ (Fig 4B).

Since the ‘Red’ subpopulation was the most distinct from other subpopulations; the pair-wise comparisons were made to tabulate DEGs between ‘Orange’, ‘Green’, ‘Blue’ with ‘Red’ subpopulations. This showed differentially expression of 568 upregulated & 582 downregulated genes between ‘Red’ & ‘Orange’ subpopulations; 812 upregulated & 908 downregulated genes between ‘Red’ & ‘Green’ subpopulations and 959 upregulated & 968 downregulated genes between ‘Red’ & ‘Blue’ subpopulations with log fold change of more than 2 and adjusted *p* value of less than 0.05 (Fig 4C, Supplementary Table 1). Differentially expressed genes from pairwise comparisons are also depicted as volcano plots (Supplementary Fig 11). Interestingly, ALDH^High^ (‘Red’ and ‘Green’) subpopulations showed significantly higher expression of ALDH1A3 as compared to their respective ALDH^Low^ subpopulations, ‘Orange’ and ‘Blue’ (Supplementary Fig11). Gene set enrichment analysis (GSEA) was performed from larger list of DEGs (q<0.1, log_2_FC >0.5) between Red and Orange, Red and Green, Red and Blue subpopulations to derive biological inferences of differentially expressed genes among subpopulations. We observed depletion for GPCR-related gene set in all three subpopulations compared to the ‘Red’ subpopulation, indicating loss of GPCR-related signaling may change transcription state from ‘Red’ to other subpopulations (Supplementary Fig 12A). Intriguingly, we observed enrichment of signatures related to keratinization and cornified envelope with the DEGs in ‘Green’ compared to the ‘Red’ subpopulations (Fig 4D). Similarly, 3D-spheroid cultures of the ‘Green’ subpopulation showed an enrichment of these signatures compared to 3D-spheroid cultures generated from ‘Red’ subpopulation (Fig 4E). This suggested the onset of differentiation in ‘Green’ subpopulation, even when maintained in 3D spheroid cultures.

**Figure 4:**
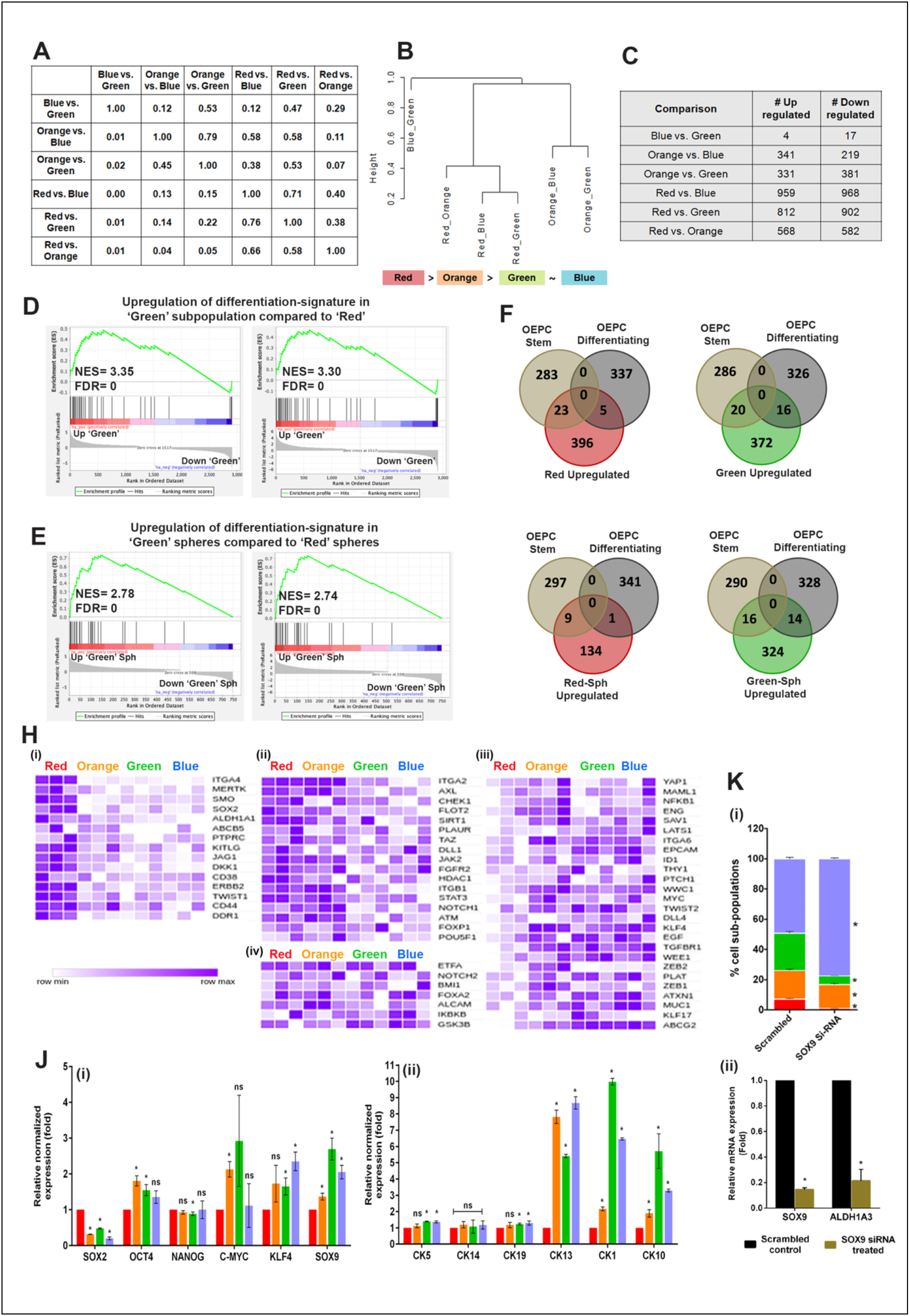
Subpopulation specific transcriptome analysis: **(A, B)** Overlap in DEGs of different pairwise comparisons. Proportion of overlap between each pairwise comparison (A). Hierarchical structure of different cell sub-populations (B). **(C)** Pairwise comparisons of differentially expressed genes of four subpopulations of GBC02 cell line. **(D)** Gene set enrichment analysis (GSEA) with genes up-regulated in ‘Green’ subpopulation as compared to ‘Red’ subpopulation from GBC02 monolayer cultures showing enrichment of genes involved in Keratinization and Cornified envelope formation. **(E)** Gene set enrichment analysis (GSEA) with genes up-regulated in Green subpopulation as compared to Red subpopulation from GBC02 3D spheroids showing enrichment of genes involved in keratinization and cornified envelope formation. **(F)** Venn diagram showing overlap of basal OEPC and basal differentiating gene sets with Red vs. Green comparison upregulated genes in Red & Green sorted subpopulations monolayer cultures and **(G)** Red & Green 3D spheroids from GBC02 cell line.**(H)** Heatmaps of mean normalized expression values from GBC02 monolayer cultures for genes expressed distinctly in (i) ‘Red’ (ii) both ‘Red’ and ‘Orange’ subpopulations(iii) commonly in ‘Orange’, ‘Green’ and ‘Blue’ subpopulations and (iv) among all the four subpopulations. **(I)** qRT-PCR of the four sorted subpopulations from GBC02 monolayer cultures for various (i) stemness genes and (ii) differentiating cytokeratins. **(J)** (i) Graph showing decreased frequency in ALDH^High^ subpopulations in SCC-070 cell line upon siRNA mediated silencing of SOX9 and (ii) qRT-PCR after siRNA knockdown of SOX9 in SCC-070 cell line.

A recent study by Jones *et. al.*, 2019 reported diversity within basal layer oral epithelial progenitor cells (OEPCs) in normal oral mucosa of mouse, using single cell RNA sequencing [29]. This study has demonstrated the maintenance of stemness with onset of differentiation in subsets of basal layer keratinocytes in mouse oral mucosa [29]. Therefore; to explore similarities, we overlapped our DEGs with the gene sets of stem/progenitors and differentiating cells of basal layer keratinocytes. Interestingly, from the DEGs between ‘Red’ and ‘Green’ subpopulations; the upregulated genes in ‘Red’ subpopulation had significantly higher overlap with OEPC-stem cell cluster (Fig 4F); whereas, the upregulated genes in ‘Green’ subpopulation had almost equal number of genes overlapped to both OEPC-stem as well as differentiating cell clusters (Fig 4F). Very similar patterns were obtained with DEGs between 3D-spheroids generated from ‘Red’ and ‘Green’ subpopulations (Fig 4G). Similar to the ‘Green’ subpopulation, upregulated genes in ‘Orange’ and ‘Blue’ subpopulations as compared to the ‘Red’ also had equal overlaps with the OEPC-stem and differentiating cell clusters (Supplementary Fig 12B). Further, we explored the enrichment of transcription factors (TFs) specific to stem/progenitors and differentiating clusters from this study. Interestingly, all four subpopulations showed mosaic expression pattern of these TFs (Supplementary Fig. 12C). This prompted us to explore if our four subpopulations exhibit the signature of stem cells. Hence, we performed single sample gene set enrichment analysis (ssGSEA) utilizing the previously reported gene-sets for Adult Tissue Stem Cells (ATSC) [30]. All four subpopulations as well as 3D-spheroids of ‘Red’ and ‘Green’ subpopulations showed positive enrichment with no significant differences among them for the stemness signature (Supplementary Fig 12D). Encouraged by these results, we further evaluated expression pattern of 65 different genes which are studied for their roles in stemness maintenance in various cancer types. Interestingly, we found specific subsets of genes expressed distinctly in ‘Red’ (Fig 4H(i)); or both ‘Red’ and ‘Orange’ (Fig 4H(ii)); or commonly in ‘Orange’, ‘Green’ and ‘Blue’ subpopulations (Fig 4H(iii), or among all the four subpopulations (Fig 4H(iv)). This suggested that stemness may be maintained in these subpopulations and regulated by distinct gene expression networks.

To define the cell state specific gene expression signatures of these four subpopulations, we generated a list of unique-DEGs specific to each subpopulation (Supplementary Table 2). Since, the ‘Green’ and ‘Blue’ subpopulations showed similar gene expression, the ‘Blue’ subpopulation was excluded from this analysis. Gene Ontology (GO) analysis for subpopulation specific unique-DEGs using the cytoscape tool-BiNGO, resulted in overrepresentation of specific pathways for these subpopulations (Supplementary Table 3,4,5). Interestingly, while the ‘Red’ subpopulation with CD24^Low/^ALDH^High^ phenotype overrepresented the process of organ development and lipid metabolism, the CD24^High/^ALDH^High^, ‘Green’ subpopulation had overrepresentation of organ development with differentiation processes.

Based on these results, we inferred that the spontaneous emergence of CD24^High/^ALDH^High^ subpopulation from the CD24^Low/^ALDH^High^ is a process of cellular differentiation, retaining mixed transcriptional signatures of Intermediate states of stemness and differentiation. We validated these results in spheroids generated from the four subpopulations of GBC02 by qRT-PCR for TFs responsible for maintenance of stemness (Fig 4J-i) as well as cytokeratin markers associated with basal and differentiated suprabasal layer of keratinocytes (Fig 4J-ii). While all four subpopulations-derived spheroids maintained basal cytokeratin markers *CK14*, *CK5*, *CK19* at similar levels; ‘Green’, ‘Orange’ and ‘Blue’ subpopulations expressed differentiation markers *CK13*, *CK1* and *CK10* at significantly higher levels compared to the ‘Red’ spheroids. Interestingly, while *KLF4*, *cMYC*, *OCT4* and *NANOG* expression was variable among all subpopulations; *SOX2* was significantly downregulated while *SOX9* was expressed at higher level in ‘Green’, ‘Blue’ and ‘Orange’ spheroids compared to the ‘Red’ spheroids. siRNA mediated knockdown of *SOX9* expression in SCC-070 cells indeed displayed alteration in phenotype distribution (Fig 4K-i) and lower expression of *ALDH1A3* (Fig 4K-ii), suggesting the crucial role of *Sox9* in maintaining ALDH^High^ phenotypes.

### Evaluation of prognostic value of population-specific gene expression pattern

To explore the functional relevance of cell state transitions, we utilized unique-DEGs to test the correlation with clinical outcome of HNSCC patients from TCGA cohort. Using subpopulation specific unique-DEGs, mean ssGSEA scores were calculated for HNSCC patients from TCGA gene expression data and classified them into ‘low’ or ‘high’ groups. The survival of these groups was estimated using Kaplan–Meier (KM) curves and Cox-regression analyses. Importantly, among all the comparisons, statistically significant difference in survival was obtained with the top 50 and 80 upregulated unique DEGs in ‘Red’ subpopulation (Fig 5A-i and Supplementary Fig 14A). Conversely, patients with higher ssGSEA-score calculated from the most downregulated unique-DEGs showed a trend of poorer prognosis (Fig 5B-i and Supplementary Fig 14B). We verified this result by an alternate approach where survival analysis was performed using the most upregulated or downregulated unique-DEGs as query on cBioPortal web tool [31]. Similar to the results obtained with ssGSEA score; patients with alterations in upregulated unique-DEGs correlated significantly with better prognosis (Fig 5A-ii, iii); whereas, alterations in downregulated unique-DEGs significantly correlated with poorer progression and disease-free survival (Fig 5B-ii, iii). Since, unique DEGs were highly expressed in the respective subpopulation and showed minimal expression and variance in other subpopulations; therefore, one of the possible interpretations for this observation could be that patients with ‘Red’ subpopulation that lacked the ability to generate other subpopulations were likely to have better survival as compared to those where ‘Red’ subpopulation was able to generate heterogeneous subpopulations, resulting in higher intratumoral heterogeneity (ITH).

**Figure 5:**
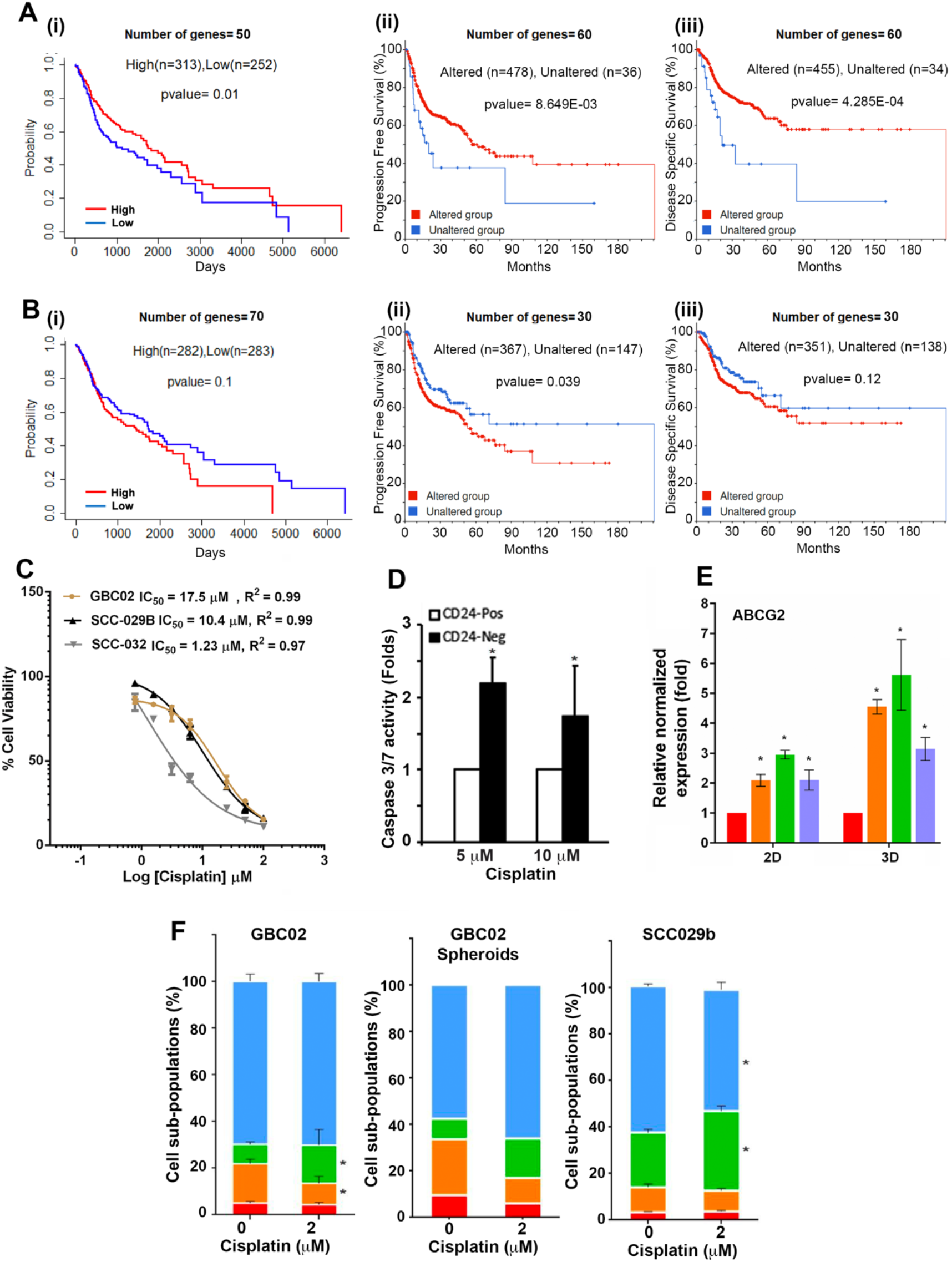
Prognostic significance of emergence of CD24^High^ Cisplatin tolerant subpopulations: (A) (i) Kaplan Meier curves of HNSCC patients segregated into High expression (Red line) and Low expression (Blue line) groups based on top 50 uniquely up-regulated genes of ‘Red’ subpopulation. (ii)Kaplan Meier curves for progression free survival and (iii) Disease specific survival for HNSCC patients having altered (Red line) or unchanged (Blue line) expression for top 60 upregulated genes in ‘Red’ subpopulation. (B) (i) Kaplan Meier curves of HNSCC patients segregated into High expression (‘Red’) and Low expression (‘Blue’) groups based on top 70 uniquely down-regulated genes of ‘Red’ subpopulation. (ii)Kaplan Meier curves for progression free survival and (iii) Disease specific survival for HNSCC patients having altered (Red-line) or unchanged (Blue-line) expression for top 30 downregulated genes in ‘Red’ subpopulation. (C) Cell survival percentages and the IC50 values of GBC02, SCC-029 and SCC-032 cell lines 3D-spheroid cultures with Cisplatin treatment. (D) Graph showing higher Caspase 3/7 activity in GBC02 cell line’s CD24^Low^ (CD24-Neg) cells compared to CD24^High^(CD24-Pos) cells upon Cisplatin treatment.(E) qRT-PCR result of ABCG2 expression in GBC02 2D sorted and 3D spheroid subpopulations. (F) (i) Graph showing significant increase in ‘Green’ subpopulation upon Cisplatin treatment for 48 hours in GBC02 cell line. Error bars represent Mean ± SEM from three biological repeats. Similar increase in ‘Green’ cells with 48 hours Cisplatin treatment in (ii) GBC02 3D spheroids (iii) SCC-029 parent cell line.

To explore this postulation, we calculated IC_50_ value of Cisplatin among three different oral cancer cell lines with distinct ‘Red’ subpopulation driven heterogeneity pattern (Fig 5C). We found that 3D-spheroid cultures of heterogeneous cell lines GBC02 and SCC-029B (where the four subpopulations originates from ‘Red’ subpopulation) exhibited higher IC_50_ of Cisplatin compared to SCC-032 cell line (where ‘Red’ subpopulation fails to generate ‘Green’ subpopulation (Supplementary Fig 13B), highlighting the importance of spontaneous generation of CD44^+Ve^/CD24^High^/ALDH^High^ ‘Green’ subpopulation from CD44^+Ve^/CD24^Low^/ALDH^High^ ‘Red’ subpopulation (Fig 5C). Next, we tested the relative Cisplatin sensitivity of CD24^High^ and CD24^Low^ subtypes of cells isolated from GBC02 cell line. Interestingly, even at higher dose (10 µM) of Cisplatin treatment for 48 Hrs, CD24^Low^ subtypes showed significantly higher Caspase 3/7 activity, indicative of higher induction of apoptosis compared to CD24^High^ subtypes of cells (Fig 5D). RT-PCR analysis revealed highest expression of drug efflux gene ABCG2 in the ‘Green’ subpopulations among all four in both 2D and 3D-spheroid culture of GBC02 in treatment naïve condition (Fig 5E), which maybe a possible explanation for Cisplatin refraction by CD24^High^ subpopulations. Further, the heterogeneous parent cultures in 2D as well as 3D-spheroid conditions, significantly enriched for CD44^+Ve^/CD24^High^/ALDH^High^ ‘Green’ subpopulation after 48 Hr of exposure to sub-lethal dose of Cisplatin (2 µM) (Fig 5F, Supplementary Fig 13A). Therefore, spontaneous transition of CD24^Low^ to CD24^High^ subtype and Cisplatin induced plasticity of ALDH^Low^ to ALDH^High^ state collectively result in emergence of CD44^+Ve^/CD24^High^/ALDH^High^ ‘Green’ subpopulation as Cisplatin tolerant state of cells, resulting in better survival of cells.

## Discussion

Phenotypic markers have been successfully employed to study ITH, biology of phenotype switching, and cancer cells survival strategies in response to therapies [19, 32, 33]. Here, using markers of putative oral-SLCCs, we have characterized diversity among CD44-positive oral cancer cells for their distinct expression of CD24 and ALDH-activity. Importantly, these characterized subpopulations served as experimental models to reliably demonstrate the spontaneous or Cisplatin driven population trajectories, and their distinct transcriptional states. We also established the clinical relevance of phenotypic cell states driven ITH in oral cancer patients. Our study strongly suggested strict unidirectional conversion of CD24^Low^ to CD24^High^ phenotype; however, stochastic bidirectional plasticity was observed on ALDH-axis in both CD24^Low^ and CD24^High^ phenotypes of oral cancer cells. Our experimental observations were validated by mathematical modelling where CD44^+Ve^/CD24^Low^/ALDH^High^ (‘Red’) subpopulation while self-sustaining itself, repopulated all other subpopulations; thus, reconstituting the phenotypic heterogeneity. Additionally, population trajectories also highlighted the sufficiency of CD24^High^ subpopulation to maintain itself in long-term 2D and 3D cultures in ‘Red’ subpopulation depleted conditions, without repopulating it. Similarly, CD44^+Ve^/CD24^High^/ALDH^Low^ (‘Blue’) subpopulations showed highest self-sustenance with the majority of transition paths leading to this subpopulation. Collectively, our observations are in concordance to the stochastic interconversions of different phenotypic subpopulations on ALDH-Axis, as reported in breast cancer cells [19, 32, 33] as well as suggest a novel strict hierarchical structure between these subpopulations on CD24-axis in oral cancer cells.

Recently, heterogeneity within mice oral mucosal basal layer has been reported [29], where in addition to harboring oral epithelial progenitor cells (OEPCs), the basal layer also accommodates maturing keratinocytes; thus revealing a continuum of cell differentiation states within it. Interestingly, some of these differentiating keratinocytes continued to co-express high levels of cytokeratin 14 (CK14) and genes associated with both OEPCs and differentiation processes, representing transitional intermediated cell states. Further, most of the basal layer cells were found to be cycling. We could draw similarities between this tissue hierarchy in normal mice oral mucosa and population trajectories in oral cancer cells. As anticipated, the ‘Red’ subpopulation with CD44^+Ve^/CD24^Low^/ALDH^High^ phenotype showed greater similarity with the transcriptome of OEPCs, supporting the reports of enrichment of putative Oral-SLCCs in CD44^+Ve^/ALDH^High^ cell population. Strikingly, other subpopulations emerging from the ‘Red’ subpopulation had transcriptome overlaps with both OEPCs and differentiating cells of oral mucosal basal layer cells, representing the transitional intermediated state of cells. The GSEA result with the differentially expressed genes between ‘Red’ and ‘Green’ subpopulation clearly showed the onset of differentiation in ‘Green’ subpopulation. Thus, the nonreversible transition of CD24^Low^ to CD24^High^ phenotype cells is possibly due to the onset of differentiation in CD24^High^cells. Also, the DEGs between ALDH^High^ (‘Red’) and ALDH^Low^ (‘Orange’ and ‘Blue’) subpopulations did not result in any specific gene sets enrichment (data not shown). This could be due to the plastic nature of ALDH^Low^ subtype of cells which may have cells with mixed transcriptome states masked in the bulk RNA-sequence data and needs exploration in future.

Further, while Sox2 was highly expressed in ‘Red’ subpopulation, other subpopulations showed downregulated Sox2 and upregulated Sox9 expression. In concordance with a recent report where depletion of Sox9 had resulted in downregulation of ALDH1 in both Sox2 and Sox9 expressing oral cancer cells [34], we too observed downregulation of ALDH1A1 expression and loss of ALDH^High^ ‘Red’ and ‘Green’ subpopulations upon Sox9 silencing . Both Sox2 and Sox9 expression are known to play important roles in maintaining cancer stemness [35, 36]. Upregulated Sox9 expression is linked with lineage infidelity driving wound repair and cancer of squamous cells [37]. Therefore, we may suggest that the ‘Green’ ‘Orange’ and ‘Blue’ subpopulations may represent intermediate transitional cell states, enriched with Sox9-positive alternate state of stemness. *In vitro* spheroid formation results from multiple independent oral cancer cells lines strongly supported this possibility and must be explored in future. To the best of our knowledge, this is the first study in oral cancer where phenotypic states of cells could be mapped to the transcriptome states demonstrating the cellular hierarchy. Moreover, our study for the first time proposed the possibility of alternate states of putative stemness maintained within intermediate differentiating subpopulations in oral cancer cells irrespective of their genetic and epigenetic diversity.

The impact of ITH on clinical outcomes is a major focus of cancer research [38, 39]. Tumor samples are usually employed to estimate the extent of ITH using genomics approaches [40, 41]. However, efforts on developing genomics-based data for investigating ITH and its clinical significance using isogenic subpopulations from patient derived cell cultures, have been limiting. Patient-derived cell cultures not only serve as essential tools for modeling disease heterogeneity but also are essential for functional validation of molecular mechanisms associated with observed complexities. Insights gained from such models can be cross-validated and translated using patient cohort. This is evident from our study, where patients with high-abundance of ‘Red’ subpopulation specific top-unique DEGs were identified to have better prognosis in HNSCC-TCGA cohort. Since, Sox2^High^/Sox9^Low^ gene expression pattern was higher in ‘Red’ subpopulation and loss of Sox2 and gain of Sox9 was the pattern in ‘Green’ ‘orange’ and ‘Blue’ subpopulation; our study showed strong concordance with a recent study which reported better prognosis in Sox2^High^/Sox9^Low^ expressing group compared to Sox2^Low^/Sox9^High^ expressing group in HNSCC-TCGA patient cohorts [34]. One of the possible mechanisms responsible for this observation could be the population dynamics described by us, where loss of ‘Red’ subpopulation specific unique-DEGs resulting in accumulation of Sox2^Low^/Sox9^High^ expressing cells in aggressive oral tumors with poor prognosis.

Tumor heterogeneity is a major contributor to therapy failure. One of the emerging possibilities is the transition of cancer cells across a continuum of states in response to chemotherapy to develop drug tolerance [42]. In this study, we have demonstrated that transient exposure to low doses of Cisplatin was sufficient to induce transition of ALDH^Low^ subpopulations (‘Orange’ and ‘Blue’) into the ALDH^High^ subpopulations (‘Red’ and ‘Green’); whereas, higher dose had resulted in increased induction of apoptosis in cells with CD24^Low^ phenotype status. With use of CD24^Low^-depleted cells, low-dose Cisplatin treatment and mathematical modelling, we validated that the accumulation of drug tolerant CD44^+Ve^/CD24^High^/ALDH^High^ subpopulation was a result of the induction of phenotype switching rather than selection by Cisplatin, in multiple oral cancer cell lines. Indeed in earlier studies, cells with CD24^High^-phenotype marked a transient chemo-resistant cell state in lung cancer [28] and breast cancer [32]. Also, increased CD24 expression with Oct4 and Nanog is recently correlated with poor chemo-radio therapy response and unfavorable prognosis in oral cancer [43]. Interestingly, Cisplatin selected oral cancer cells were shown to express higher levels of Sox9 [34]; similar to high SOX9 expression in CD44^+Ve^/CD24^High^/ALDH^High^ subpopulation in our study, too. Therefore, we support the notion that phenotype switching may act as a robust mechanism for cancer cells to rapidly acquire a drug tolerant state in changing environment from where these cells may eventually emerge as a drug-resistant population in relapsed tumor. Our cellular models may serve as tool to define the possible mechanisms underlying cross-talk among phenotypes and identify targets against the population-dynamics, as novel therapeutic strategy.

In summary, our study identified the maintenance of cell-state proportions through the process of cellular differentiation as well as spontaneous interconversions. Further, we unmasked the ability of oral cancer cells of gaining adaptive-resistance against Cisplatin by altering the inter-conversion rates of cell-states transitions *in vitro,* irrespective of their genetic and epigenetic diversity. With this report we are highlighting our ability to relate cell-states for theoretical and experimental studies and its translational implications to the clinics. Our study provides the prerequisite knowledge which defines the composition of subpopulations critical for global tumor behaviour in oral cancer, majorly lacking for most of the solid tumors. Thus, characteristics of these phenotypic subgroups may be optimum for estimating the intratumoral heterogeneity in oral cancer patients.

## Acknowledgement

This work is supported by the grant (IA/I/13/1/500908) received from Wellcome trust-DBT India Alliance. KV acknowledges DST-INSPIRE, SG acknowledges DBT and AKP acknowledges ICMR for fellowship support. We thank Professors Saumitra Das and Partha P Majumder and Dr. Kartiki Desai for scientific discussions and Dr. Arindam Maitra for assistance with transcriptomics platform. We also thank Ms. Bhaswati Tarafdar for her support and assistance with flow cytometry.

---

**Supplementary Fig. 1:**
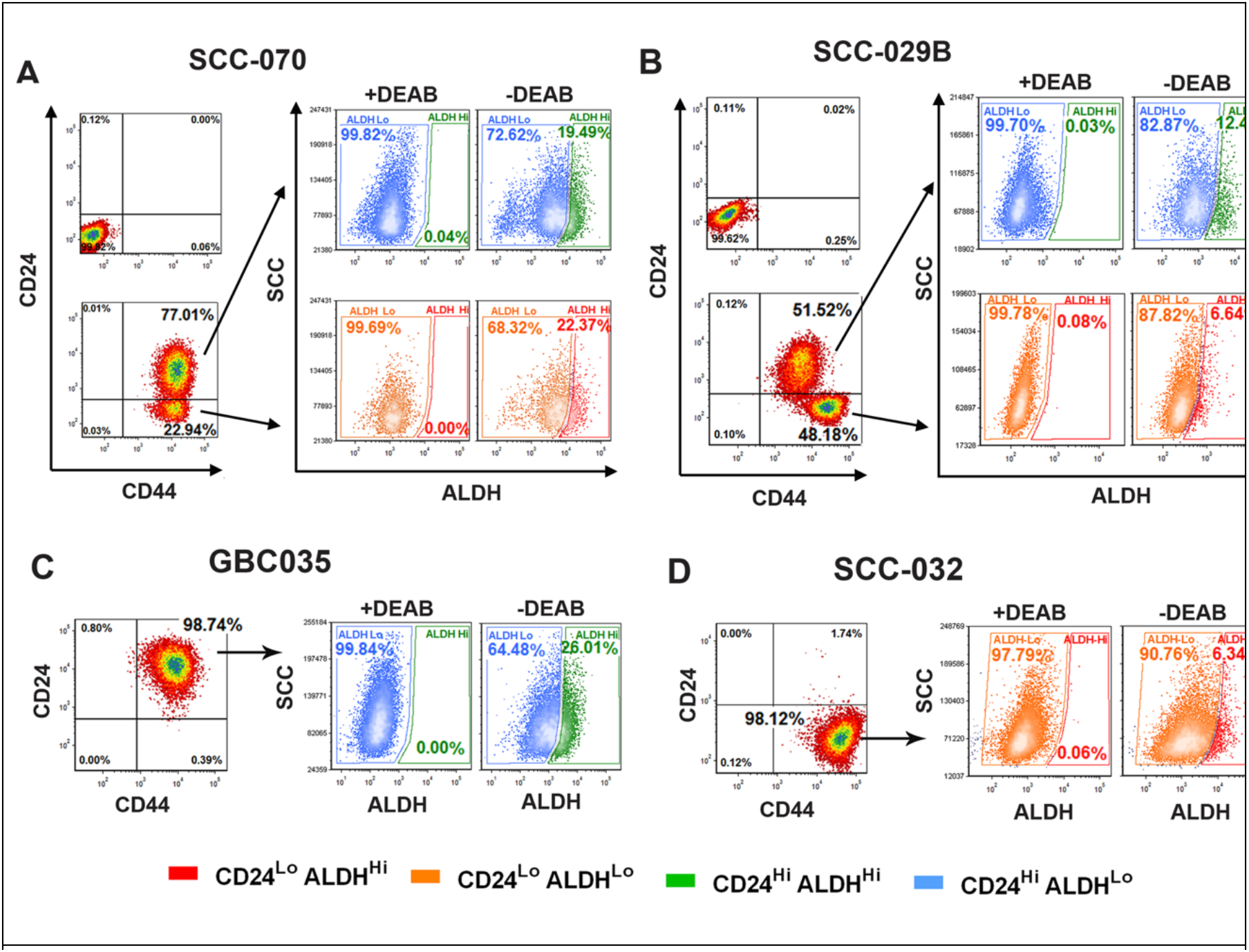
Representative FACS dot plots of **A)** SCC070 **B)** SCC029B **C)** GBC035 **D)** SCC032 cell lines CD24/CD44 staining (Left bottom) with respect to Isotype control (Left top). ALDEFLUOR (ALDH) phenotype of CD24^Hi^/CD44^+^ subpopulation (Right top) and CD24^Lo^/CD44^+^ subpopulation (Right bottom) in presence or absence of DEAB.

**Supplementary Fig. 2:**
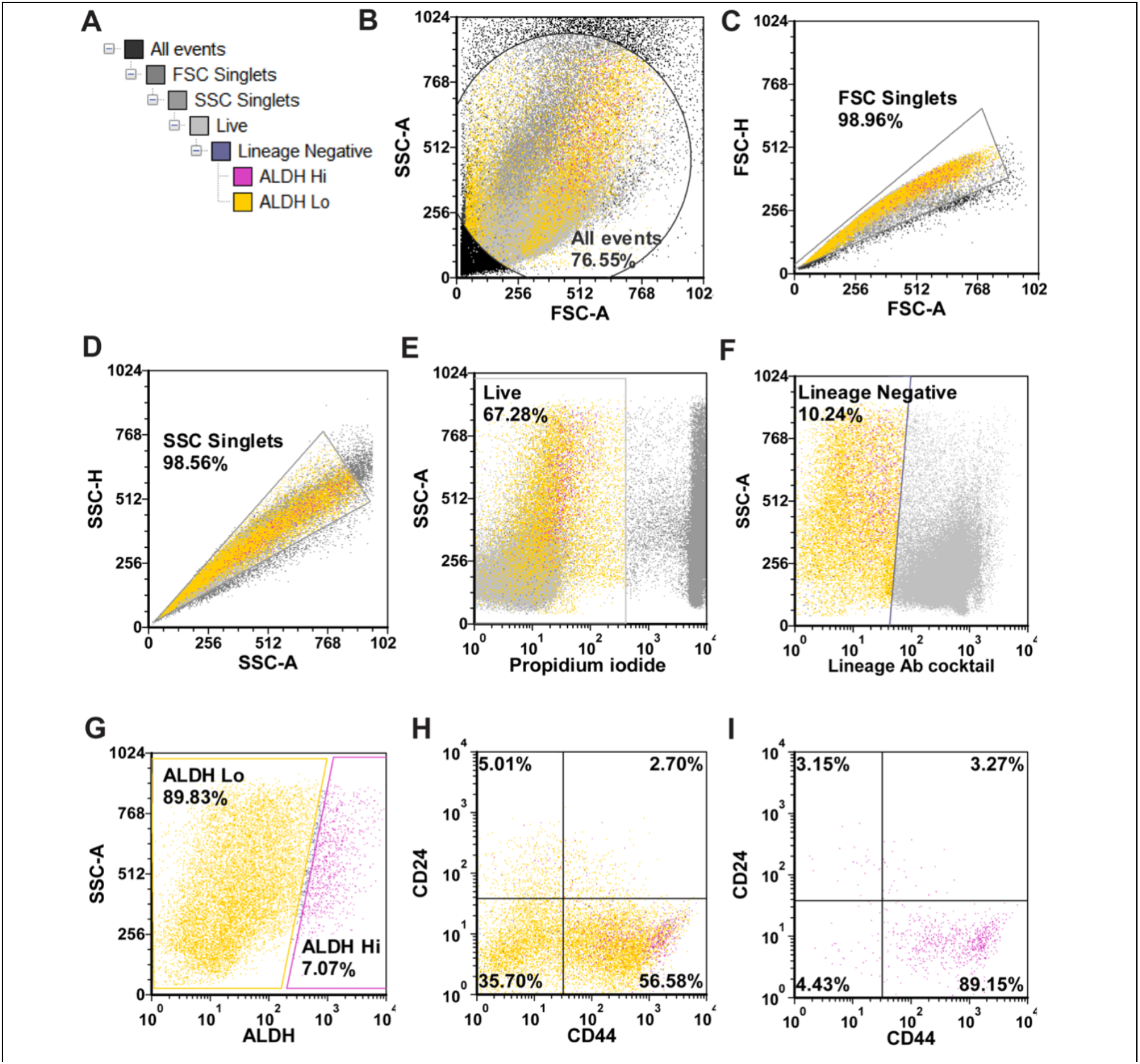
**A)** Gating strategy of ALDEFLUOR assay combined with CD24/CD44 staining in a freshly resected and dissociated patient tumor. **B)** Dot plot showing all events. **C, D)** Dot plots showing single cell selection upon doublets elimination from analysis. **E)** Selection of propidium iodide negative live cells. **F)** Selection of hematopoietic lineage negative cells using Lineage antibody cocktail. **G)** All the hematopoietic lineage negative cells segregated into ALDH^Hi^ and ALDH^Lo^ cells. **H)** CD24/CD44 phenotype of all the lineage negative cells. **I)**CD24/CD44 phenotype of ALDH^Hi^ cells only.

**Supplementary Fig. 3:**
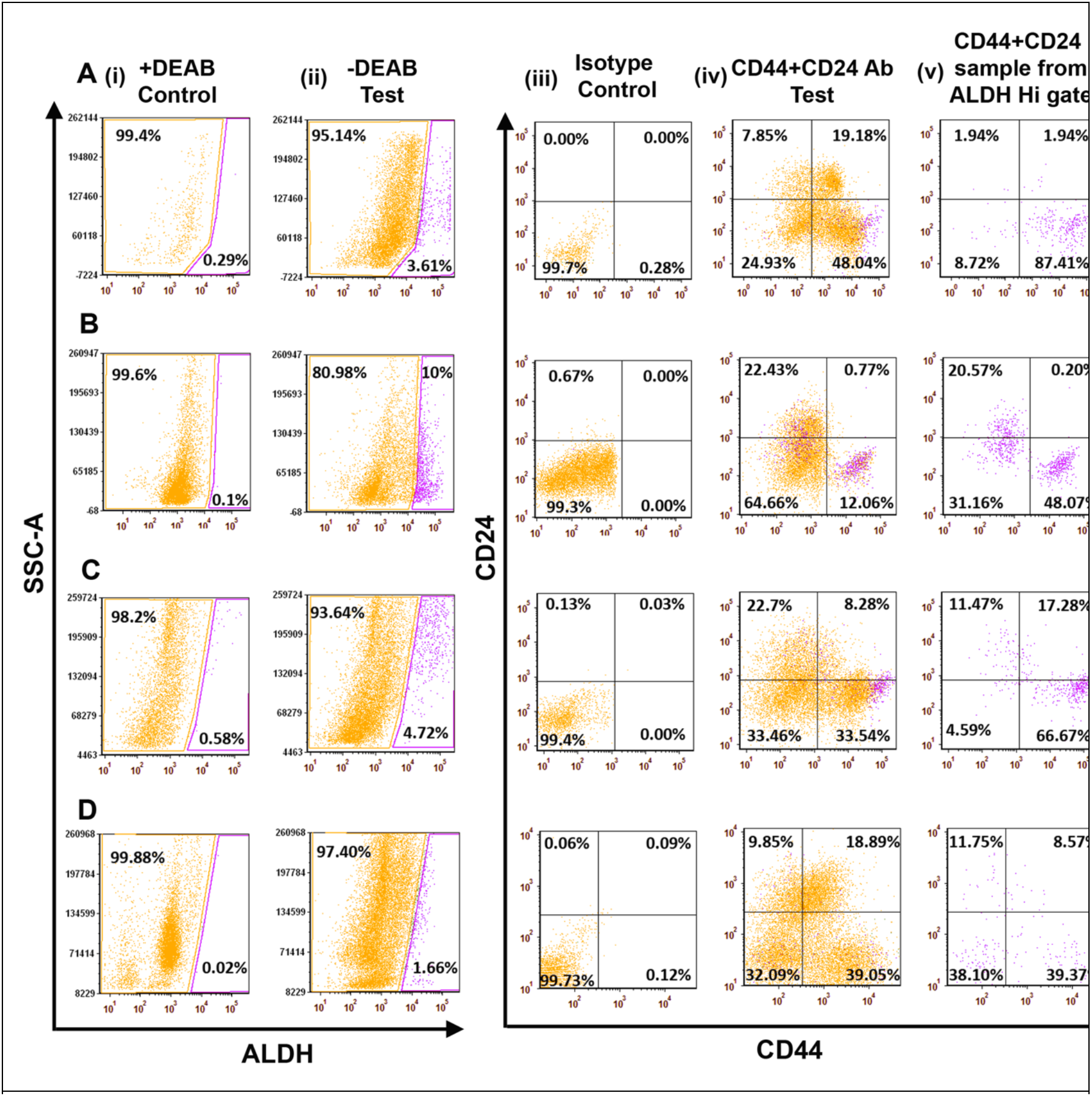
Representative dot plots of **A.** KV017, **B.** AP039, **C.** AP037, **D.** AP034 patient samples (row wise). **i)** With DEAB Control sample for setting background fluorescence for ALDH Hi cells. **ii)** Without DEAB Test sample showing ALDH^Hi^ (yellow) and ALDH^Lo^ (pink) cells. **iii)** Control sample with isotype antibodies. **iv)** Test sample with CD24, CD44 antibodies. **v)**CD24/CD44 phenotype of ALDH^Hi^ cells only.

**Supplementary Fig. 4:**
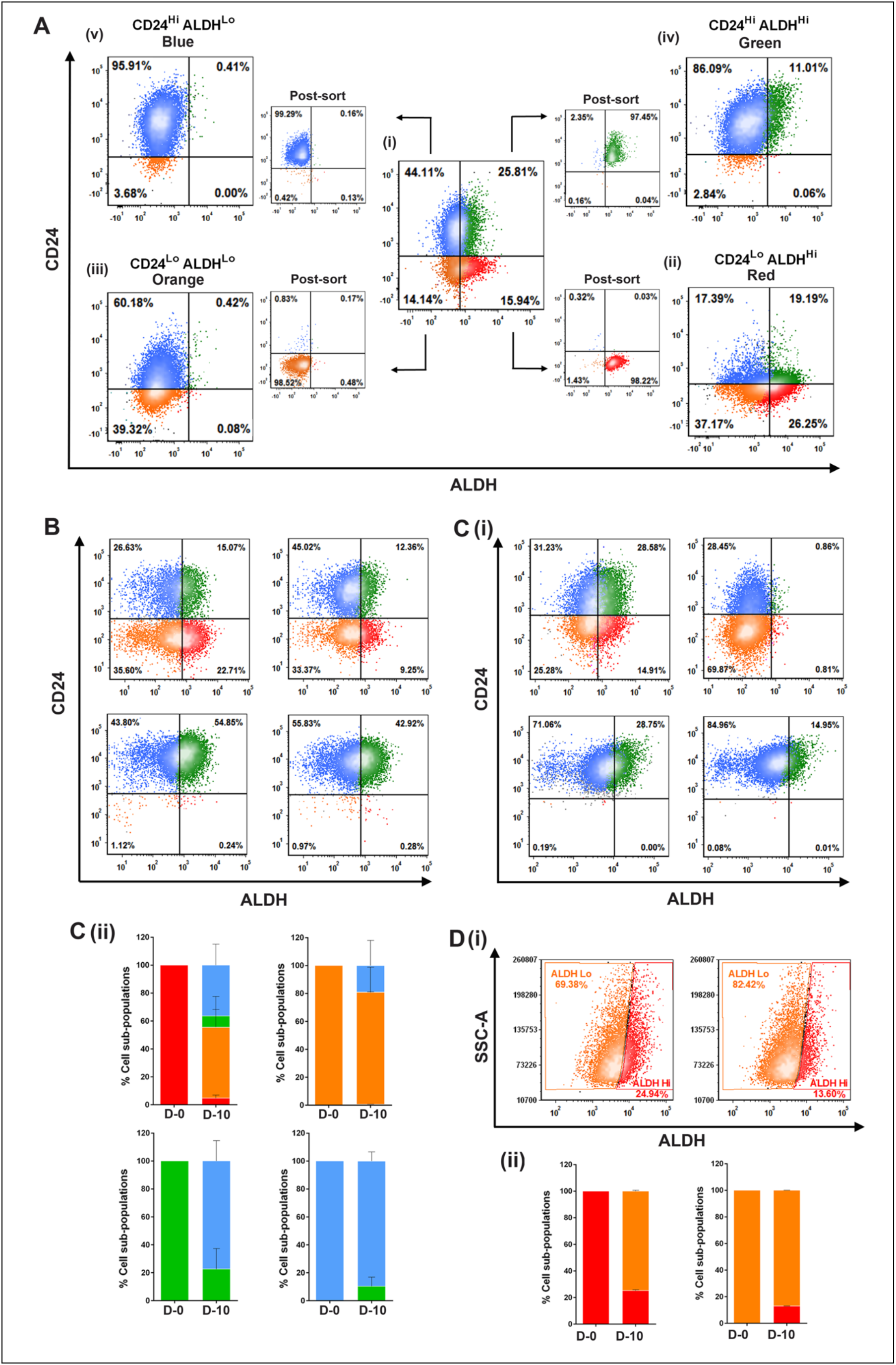
**A) (i)** Representative FACS dot plots of the GBC02 parent culture showing its CD24 and ALDH phenotype with existence of four populations and their post-sort data. Re-analysis for CD24 and ALDH showing repopulation of the **(ii)** Red **(iii)** Orange **(iv)** Green and **(v)** Blue phenotypes on Day -10 of sorting. **B)** Representative FACS dot plots of the SCC070 cell line’s Red, Orange, Green and Blue subpopulations repopulation pattern on Day-10 of sorting. **C) (i)**Representative FACS dot plots of the SCC029b cell line’s Red, Orange, Green and Blue subpopulations repopulation pattern on Day-10 of sorting. **(ii)** Repopulation frequencies of each subpopulation on Day-0 and Day-10 of sorting. Error bars represent mean ± SEM from four biological repeats. **D) (i)** Representative FACS dot plots of the SCC032 cell line’s Red and Orange subpopulations repopulation pattern on Day-10 of sorting. **(ii)** Repopulation frequencies of each subpopulation on Day-0 and Day-10 of sorting. Error bars represent mean ± SEM from two biological repeats.

**Supplementary Fig. 5:**
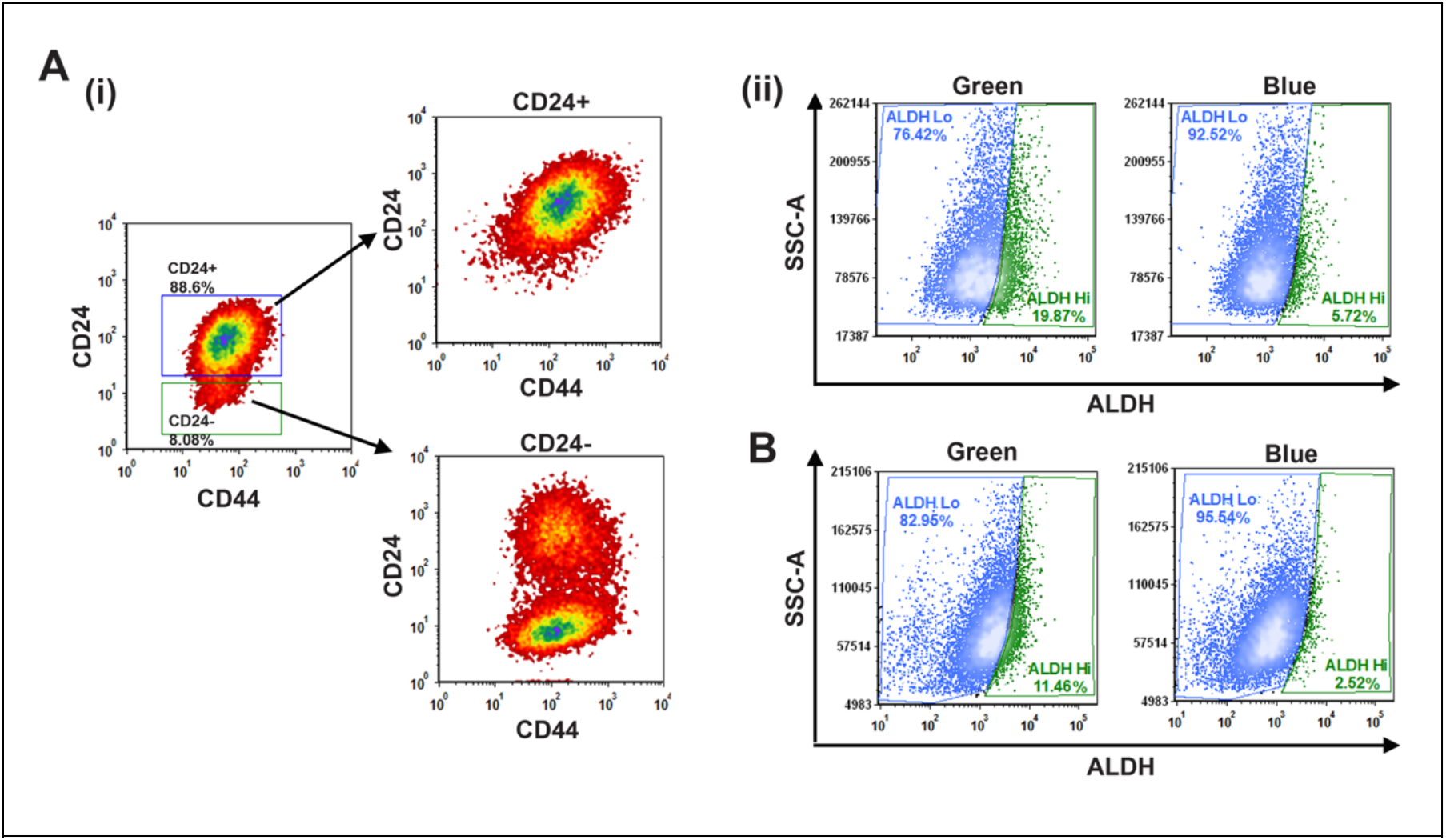
**A) (i)** Sorting of CD24+ and CD24-cells from SCC070 cell line. Established subline of CD24+ cells did not repopulate CD24-cells whereas CD24-cells repopulated both the subtypes. **(ii)** ALDH High and Low repopulation of SCC070 CD24+ subline’s sorted cells. **B)** ALDH High and Low repopulation of GBC035 sorted cells.

**Supplementary Fig. 6:**
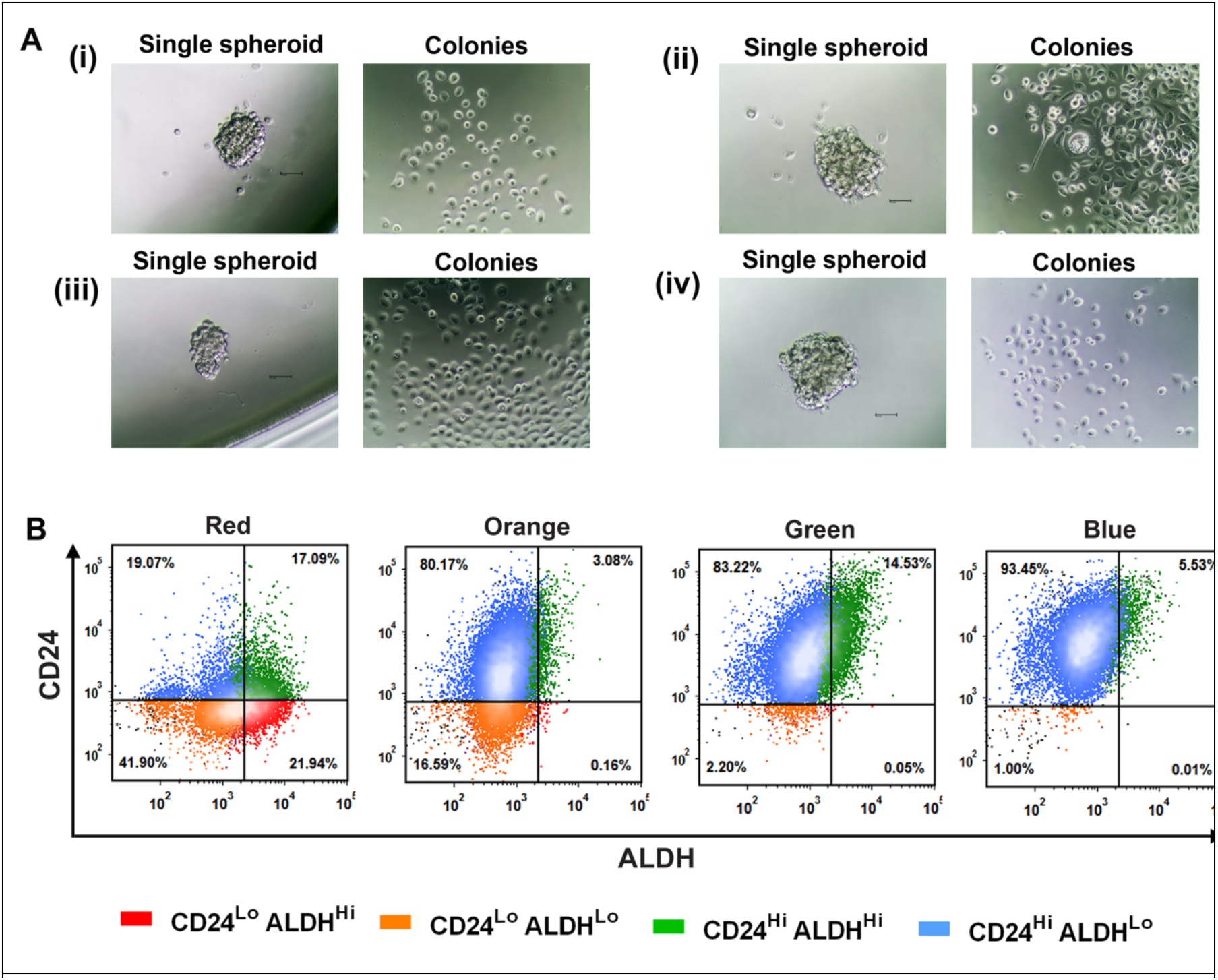
**A)** Representative pictures of single spheroids and colonies generated from them in (i) Red (ii) Orange (iii) Green and (iv) Blue subpopulations of GBC02 cell line. **B)** Representative FACS dot plots of repopulation of single spheroids of Red, Orange, Green and Blue subpopulations.

**Supplementary Fig. 7:**
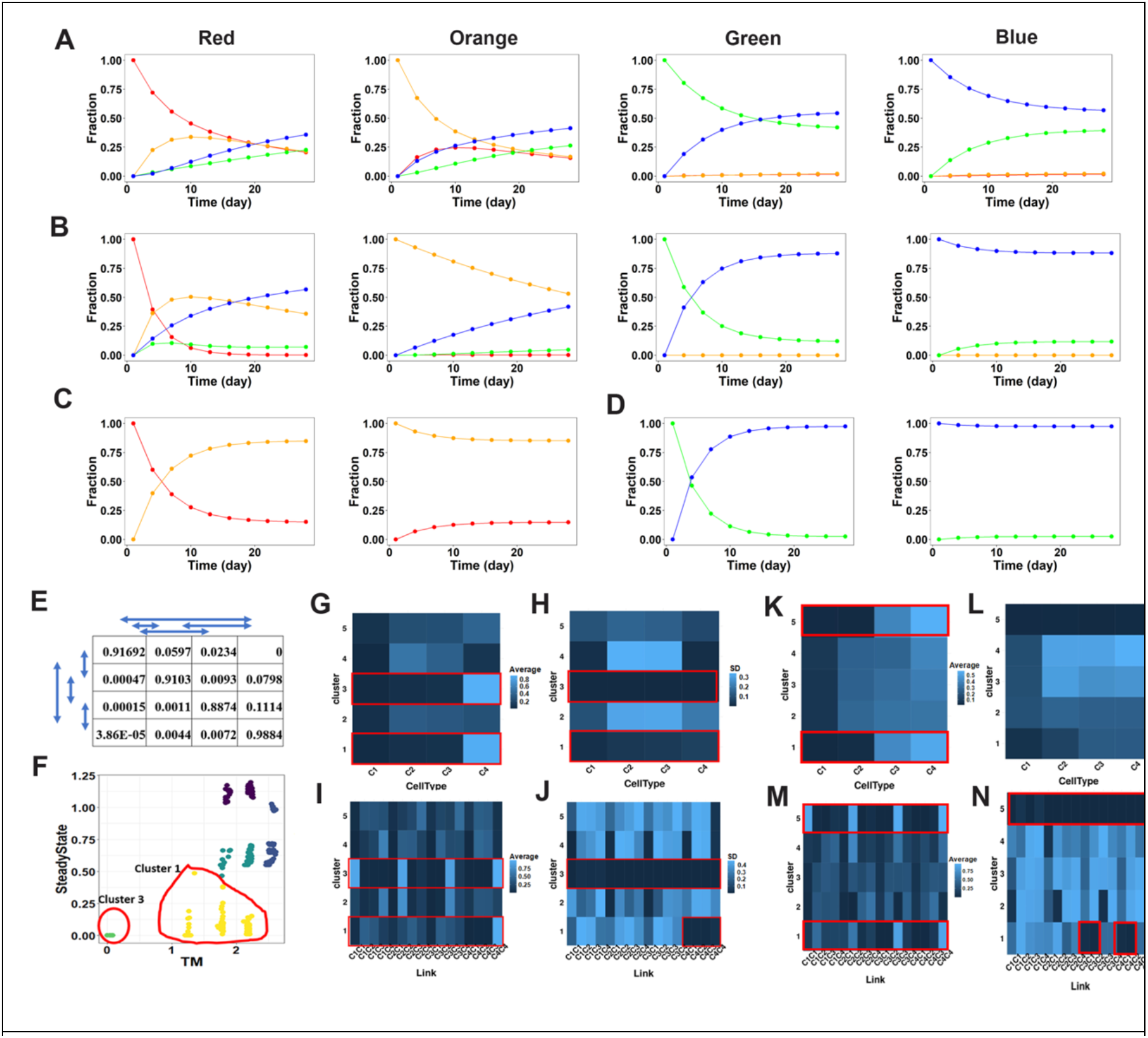
The phenotypic transition trajectories of Red, Orange, Green and Blue subpopulations of **A)** SCC070 **B)** SCC029B **C)** SCC032 and **D)** GBC035 cell lines.**E)**Demonstration of randomization of transition matrix for GBC02. **F)**Clustered scatterplot of distance of randomized transition matrices (x-axis) and the corresponding steady states (y-axis)from their wild-type (WT) counterparts, for GBC02. **G)** Cluster-wise heatmap for GBC02 of mean steady state values corresponding to the randomized transition matrices (TMs). TheCellTypes are labelled as follows: C1 – CD24^Low^/ALDH^High^, C2 - CD24^Low^/ALDH^Low^, C3 –CD24^High^/ALDH^High^, C4 – CD24^Low^/ALDH^Low^ . The highlighted rows indicate the cluster of TMs with similar steady state as WT. **H)** Same as G, but for Standard deviation instead of mean. **I)**Same as G, but for the elements of TM. The x-axis labels are to be read as follows:CiCj =Transition rate of CellType Ci to CellType Cj, i,j = {1,2,3,4}. **J)** Same as I, but for standard deviation instead of mean. Low SD regions are highlighted in red. **K)** Cluster-wise heatmaps for SCC070 cell line showing the mean and **L)** standard deviation of the steady state fractions of the CellTypes (x-axis). Labelling of the cell types is the same as in GBC02 (G-H). **M, N)** Same as (K-L) but for transition matrices of SCC070.

**Supplementary Fig. 8:**
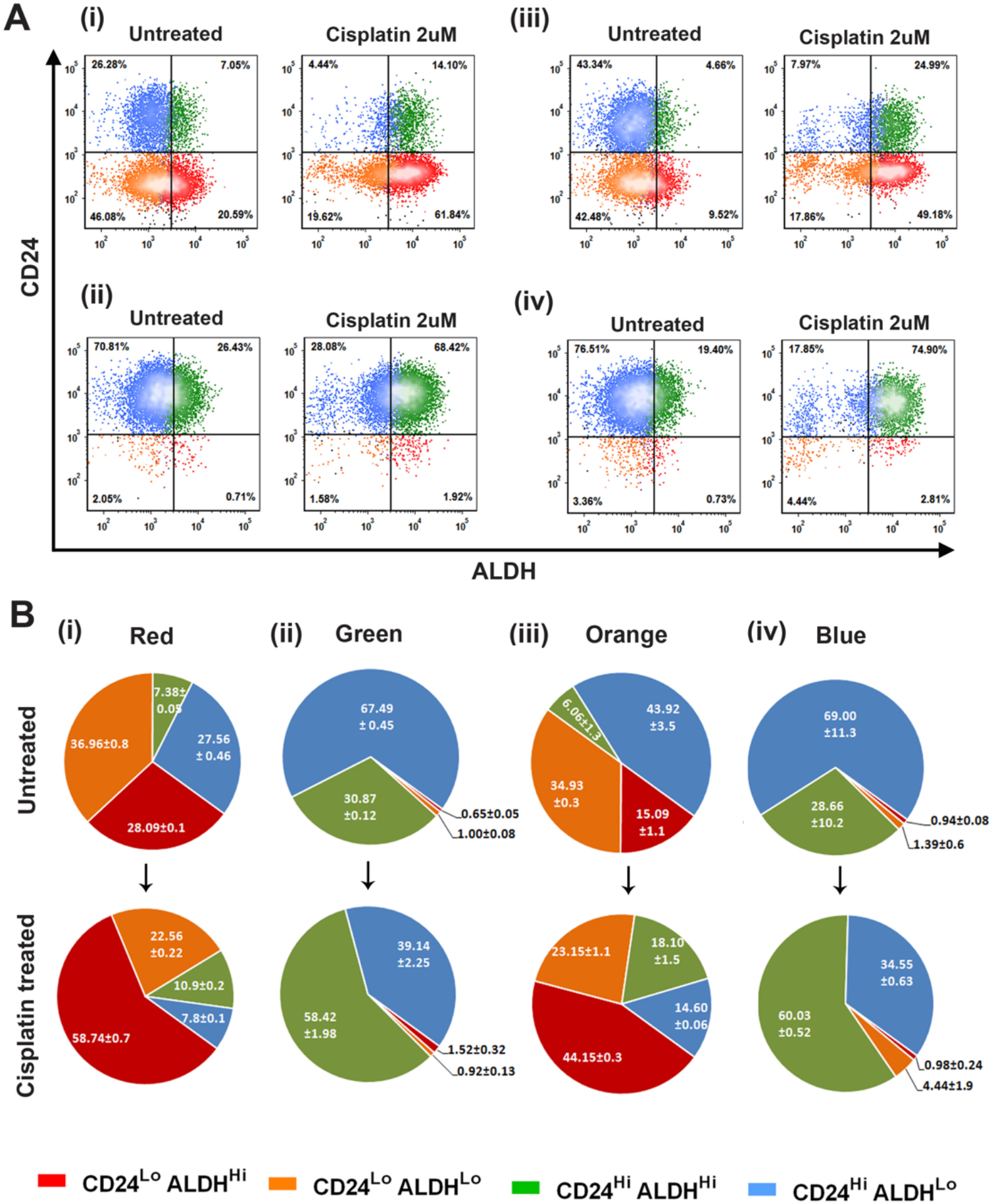
**A)** Representative dot plots of repopulation dynamics of SCC070 four subpopulations; (i) Red (ii) Green (iii) Orange and (iv) Blue in Untreated and Cisplatin (2µM) treated conditions. **B)** Repopulation frequencies of the four subpopulations (i) Red (ii) Green (iii) Orange and (iv) Blue in Untreated vs. Cisplatin (2µM) treated conditions from two biological repeats.

**Supplementary Fig. 9:**
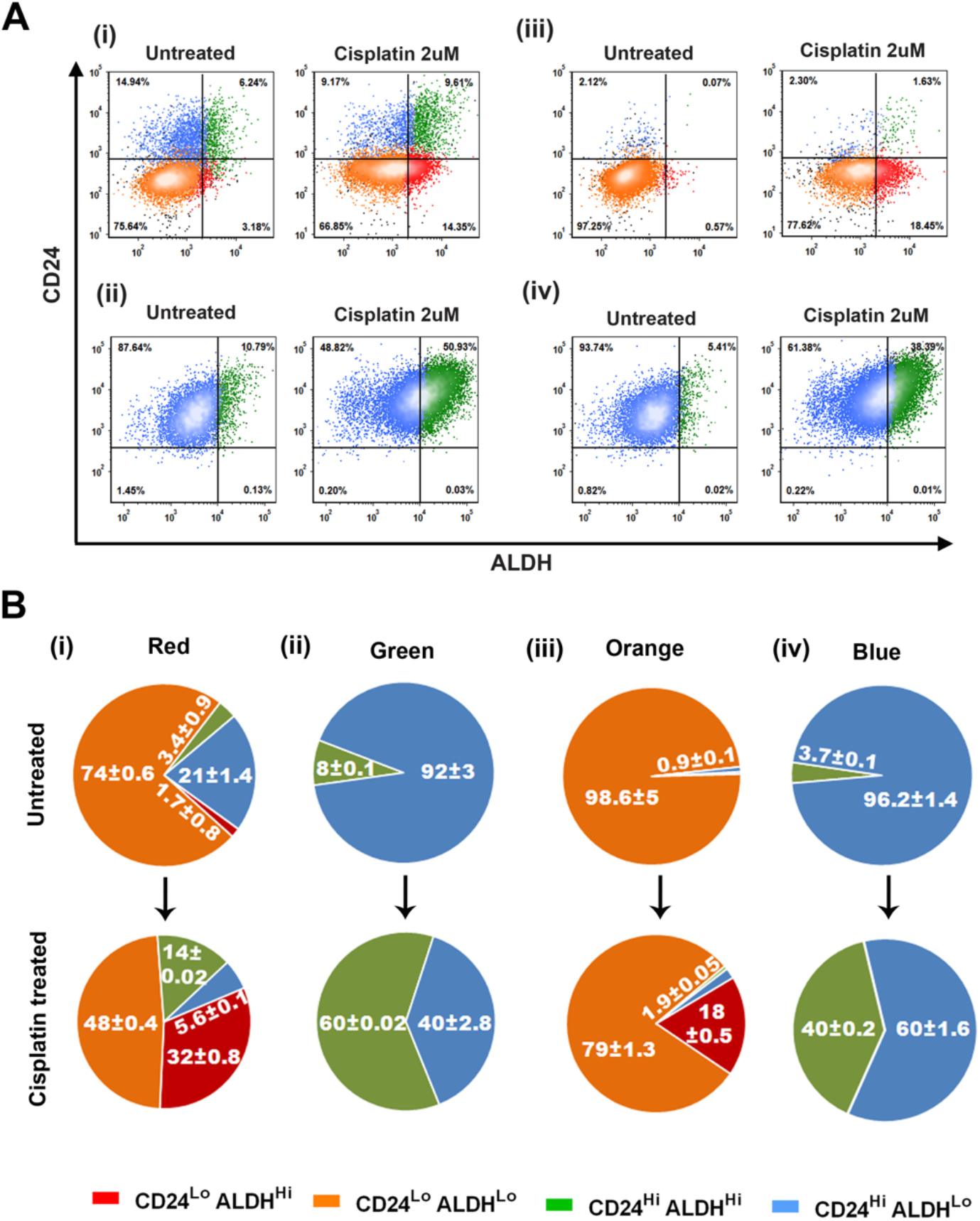
**A)** Representative dot plots of repopulation dynamics of SCC029 four subpopulations; (i) Red (ii) Green (iii) Orange and (iv) Blue in Untreated and Cisplatin (2µM) treated conditions. **B)** Repopulation frequencies of the four subpopulations (i) Red (ii) Green (iii) Orange and (iv) Blue in Untreated vs. Cisplatin (2µM) treated conditions from two biological repeats.

**Supplementary Fig. 10:**
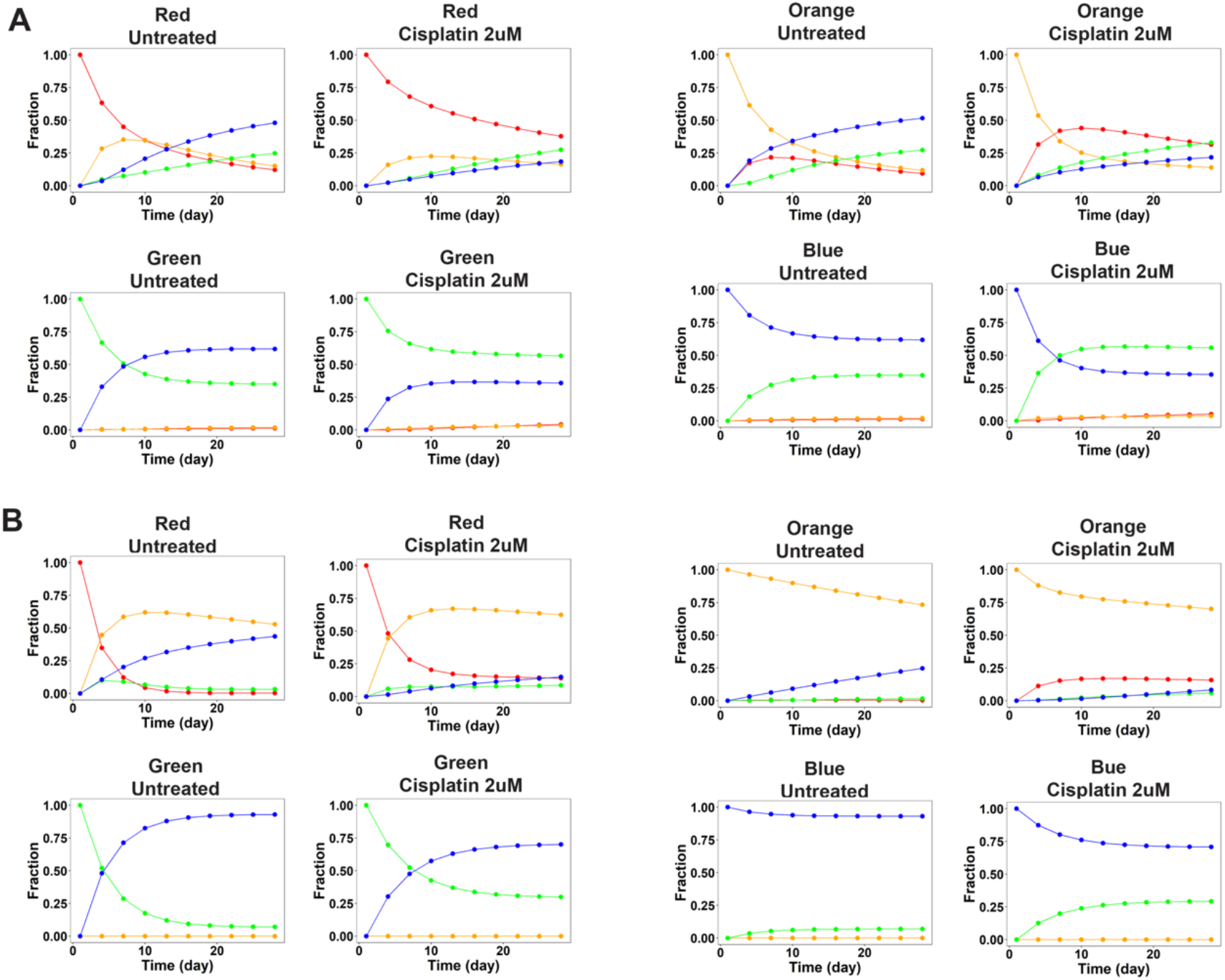
The phenotypic transition trajectories of the four subpopulations in Untreated vs. Cisplatin (2uM) treated conditions in **A)** SCC070 and **B)** SCC029B cell lines.

**Supplementary Fig. 11:**
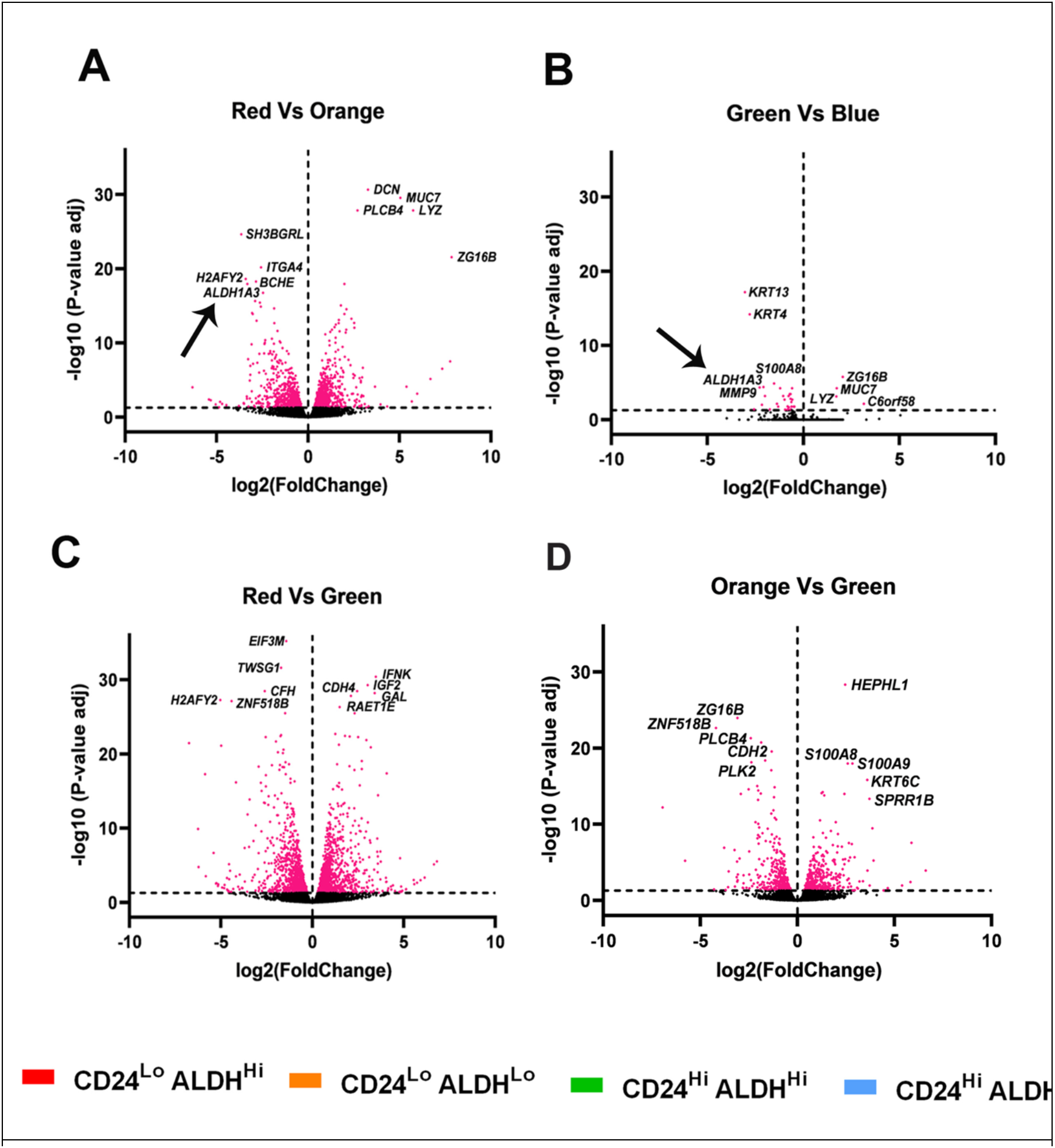
Volcano plots of **A)** Red vs. Orange, **B)** Green vs. Blue,**C)** Red vs. Green and **D)**Orange vs. Green subpopulations pair-wise comparisons. Significantly differentially expressed genes are represented in pink color. A horizontal line is used to mark the adjusted P-value threshold ≥ 0.05. The arrows point to *ALDH1A3* gene down regulated in ALDH^Lo^ subpopulations compared to their respective ALDH^Hi^ subpopulations in figures A and B. Five up regulated (right) and down regulated (left) genes are shown in each plot.

**Supplementary Fig. 12:**
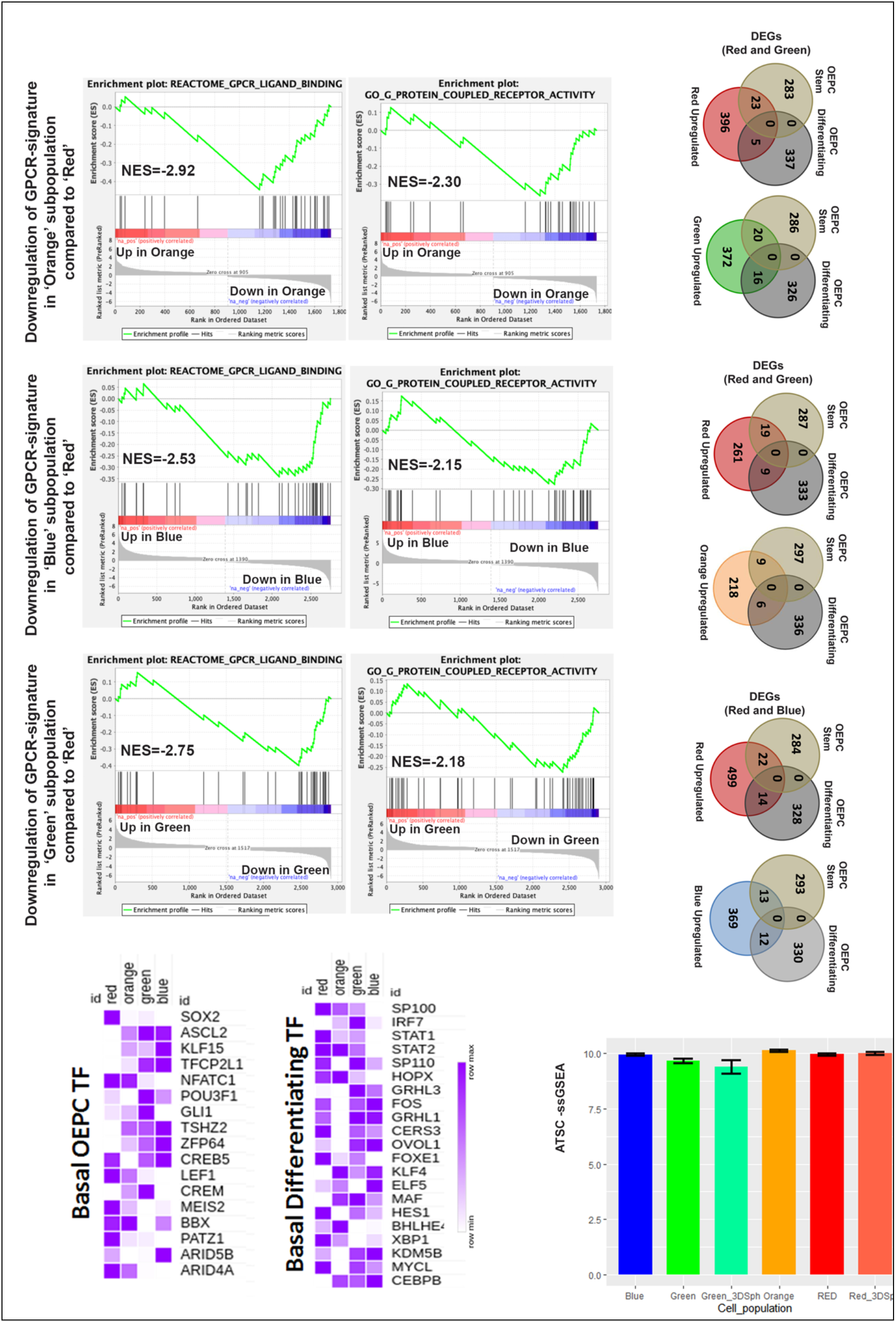
**A)** Gene set enrichment analysis (GSEA) in (i) Orange(ii) Blue and (iii) Green subpopulations compared to Red subpopulation from GBC02 monolayer cultures showing depletion of GPCR ligand binding and receptor activity. **B)** Venn diagrams showing overlap of basal OEPC and basal differentiating gene sets (Jones *et al*, 2019) with upregulated genes from **(i)** Red vs. Orange comparisons in Red & Orange sorted subpopulations, and **(ii)** Red vs. Blue comparisons in Red & Blue sorted subpopulations, from GBC02 monolayer cultures. **C)** Heatmaps of mean normalized expression values from GBC02 monolayer cultures for (i) Basal OEPC TFs and (ii) Basal differentiating TFs adapted from Jones *et al*, 2019. **D)** ssGSEA scores of adult tissue stem cell (ATSC) gene signatures in the four subpopulations ofGBC02 2D monolayers and 3D spheroids.

**Supplementary Fig. 13:**
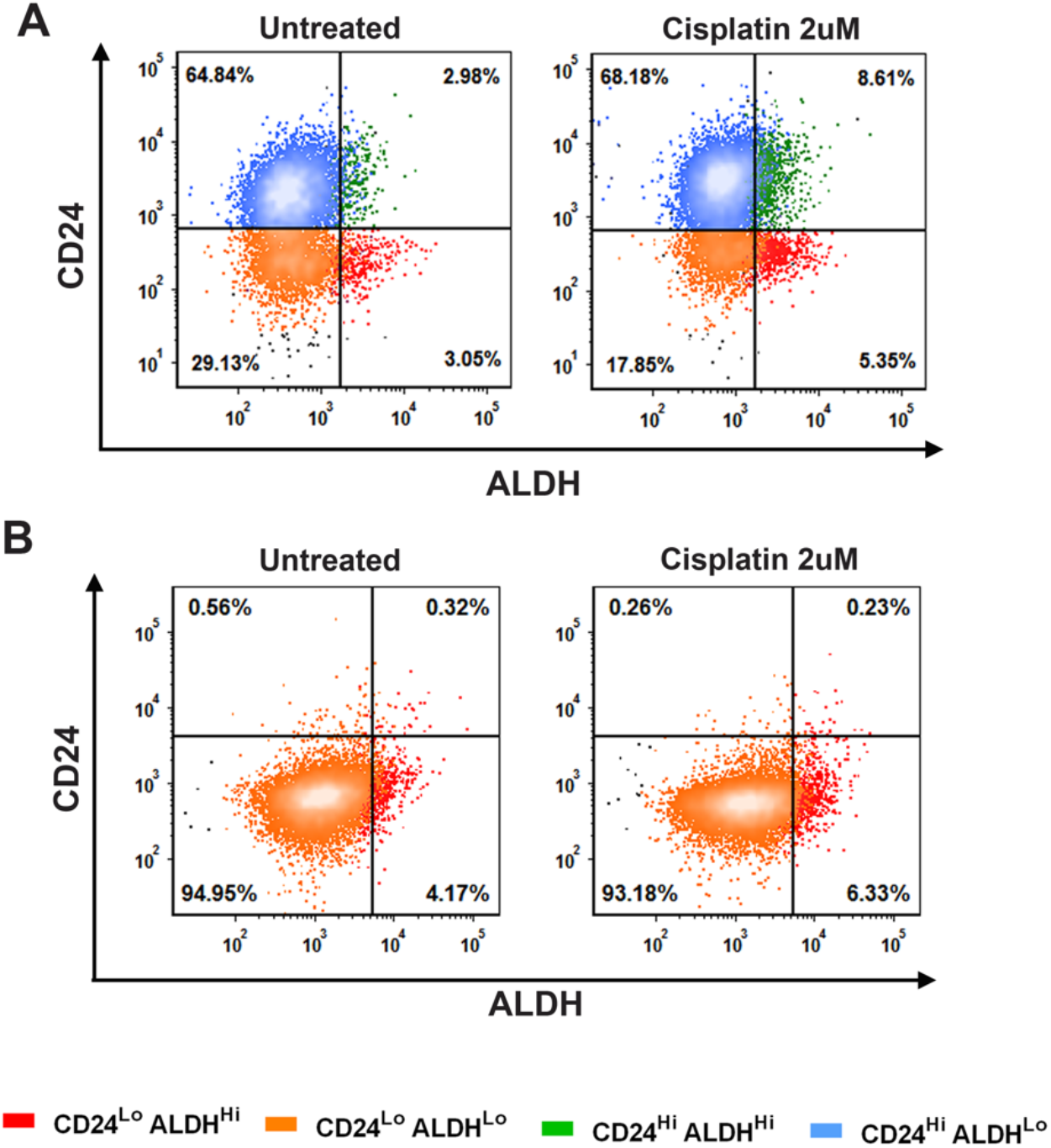
**A)** Representative dot plots of CD24 (y-axis) and ALDH (x-axis) phenotyping of GBC02 parent cell line in Untreated vs. Cisplatin (2uM) treated conditions. **B)** Representative dot plots of CD24 (y-axis) and ALDH (x-axis) phenotyping SCC032 parent cell line in Untreated vs. Cisplatin (2uM) treated conditions.

**Supplementary Fig. 14:**
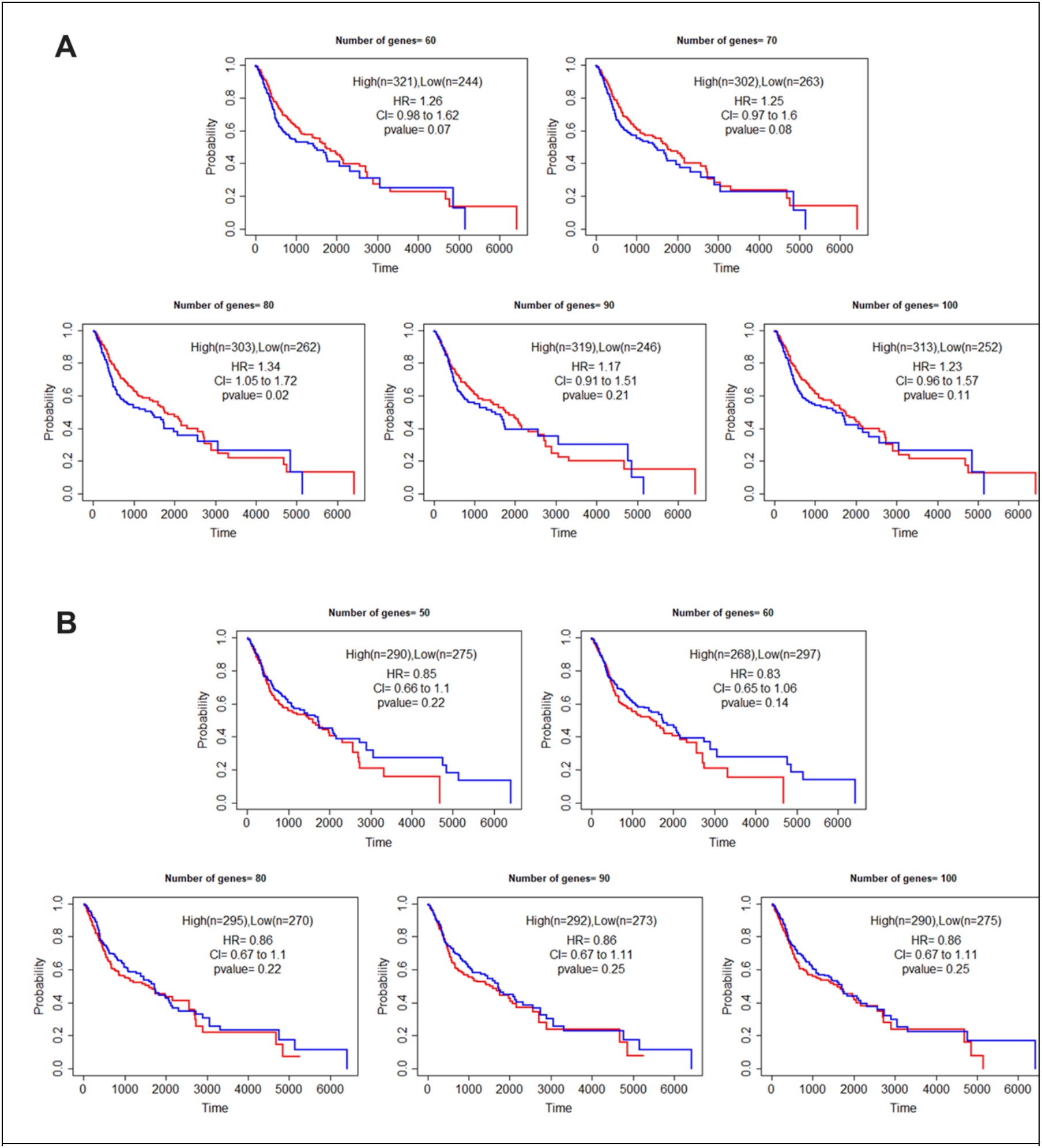
**A)** Kaplan Meier curves of HNSC patients segregated into High expression (Red) and Low expression (Blue) groups based on top 60-100 uniquely up-regulated genes of Red subpopulation. **B)** Kaplan Meier curves of HNSC patients segregated into High expression (Red) and Low expression (Blue) groups based on top 50-100 uniquely down-regulated genes of Red subpopulation.

## Notes

**Conflict of interest:** The authors declare no potential conflicts of interest.

### Competing Interest Statement

The authors have declared no competing interest.

### Summary of Updates

In this manuscript, using markers of putative oral-stem-like cancer cells (oral-SLCCs), we have characterized diversity and evolutionary dynamics among oral cancer cells. To emphasize, this is the first report in oral cancer where phenotypically distinct cells could be mapped to their transcriptome states and demonstrating the possibility of maintenance of alternate states of putative oral-SLCCs within differentiating subpopulations. Importantly, our characterized subpopulations served as experimental models to reliably demonstrate the spontaneous or Cisplatin driven population trajectories and explained by a mathematical model. Further, driven by the population dynamics; the distinct transcriptional states of diverse subpopulations were associated with clinical relevance in oral cancer patients. Importantly, this study provides the definition of the composition of subpopulations critical for global tumor behaviour in oral cancer, which is a prerequisite knowledge in precision treatment and largely lacking for most solid tumors.

